# A probabilistic and phylogenetic principal component analysis for modelling high-dimensional trait evolution

**DOI:** 10.64898/2026.05.27.728209

**Authors:** Paola Montoya, Julien Joseph, Anjali Goswami, Hélène Morlon, Julien Clavel

## Abstract

Given the ever-increasing availability of highly detailed phenotypes, modelling trait evolution in a multivariate framework is becoming a challenging task. Current phylogenetic comparative methods often struggle with high-dimensional datasets because they suffer computational limitations and interpretability. Here, we propose a maximum likelihood-based approach called Probabilistic and Phylogenetic Principal Components Analysis (P3CA) to circumvent current limitations. This approach is based on a continuous latent variable model, whereby observed traits are explained by a smaller number of unobserved variables that evolve according to a given evolutionary model. We implement the approach under Pagel’s lambda model using an Expectation-Maximisation algorithm that makes it computationally efficient and allows missing values. Using simulations, we demonstrate that evolutionary parameters are accurately estimated, regardless of phylogenetic signal, the number of traits or the proportion of missing values. The reconstruction of the reduced space is more accurate than the one obtained using other dimensionality reduction approaches, such as phylogenetic and conventional PCA. Likewise, the estimated values for missing data are more accurate than those obtained using current phylogenetic data imputation approaches. We illustrate the approach on a 3D geometric morphometric dataset describing Crocodyliformes skull shapes and containing around 4% of missing data. Our P3CA method unlocks the possibility to analyse and more easily interpret the large-scale multivariate datasets generated in recent decades within a phylogenetic comparative framework.

---

Large datasets describing high-detailed phenotypes across broad taxonomical scales, such as whole anatomical structures described from 3D-images (e.g., Phenome10k (Goswami 2015): MorphoSource (morphosource.org); MorphoMuseuM (Lebrun and Orliac 2016), DigiMorph (digimorph.org)), or complete gene expression profiles (e.g., Bgee (Bastian et al. 2025)), are becoming increasingly available. High-throughput phenotyping provides a valuable framework for studying the evolution of quantitative phenotypes, however the tools required for processing and analysing them still have limitations (Goswami et al. 2019; Snead and Clark 2022; Goswami and Clavel 2025). Current phylogenetic comparative approaches for studying phenotypic evolution in a strict multivariate context (i.e., taking into account the covariance among the traits), involve expensive matrix operations that scale with dimensions. Moreover, additional challenges arise when the datasets are high-dimensional (i.e., when the number of traits exceeds the number of species, *n* < *p*) or when the traits are highly collinear by construction (e.g., geometric morphometric data processed by Procrustes superimposition techniques where each species are scaled, rotated and translated onto a common conformation), because this prevents the use of conventional likelihood-based techniques. Although some methods have recently been developed to tackle most of these issues (e.g., (Clavel et al. 2019; Montoya et al. 2026)), they remain computationally prohibitive when applied on datasets containing more than a few thousand traits, since they rely on manipulating matrices as large as the number of traits. They also often lack interpretability due to the large number of parameters that need to be estimated (for instance, the trait variances and covariances) in order to describe the coevolution of high-dimensional traits.

An alternative for modelling trait evolution directly through high-dimensional multivariate diffusion processes (such as Brownian motion) is to reduce the dataset to fewer dimensions that can then be analysed by conventional (low-dimensional or univariate) phylogenetic comparative methods. Principal Component Analysis (PCA) is probably the most popular technique for dimensionality reduction. The PCA and its phylogenetic counterpart (phyPCA, (Revell 2009)) project the data onto independent components (or axes), from which a smaller set explaining most of the variance can be retained (see (Jolliffe 2002) for an extended explanation on the PCA). The phyPCA differs from the PCA in that it projects the main evolutionary changes – as expected under a given evolutionary model (e.g., Brownian motion) – on these components (see (Revell 2009; Polly et al. 2013)). In practice, the phyPCA is mostly limited to the Brownian motion model in high-dimensional datasets because it is the only model for which the parameters can be estimated using analytical solutions ((Revell 2009) but see (Clavel et al. 2019)). Hence, researchers often reduce datasets to a set of variables using phyPCA under the Brownian motion model before fitting other evolutionary models, which can lead to spurious results in the downstream analyses, such as an increased bias in model selection (Uyeda et al. 2015) and high type I error rates in statistical tests (Clavel and Morlon 2020).

In order to overcome these limitations, we propose a phylogenetic version of the probabilistic PCA (PPCA, (Roweis 1997; Tipping and Bishop 1999)). The probabilistic PCA is a continuous latent variable model that links the observed traits to a smaller number of latent variables forming a reduced space. Unlike PCA and phyPCA that use a deterministic approach based on linear transformations to project the data on a lower dimensional space, the PPCA assumes a probabilistic model to reduce the dimensionality. This probabilistic formulation of the PCA is convenient for extension to the case of phylogenetically structured traits, as it allows to model trait evolution directly onto the reduced space spanned by the latent variables. Another major advantage of the probabilistic PCA is the possibility to use an expectation-maximisation (EM) algorithm (Roweis 1997; Tipping and Bishop 1999; Porta et al. 2005; Li et al. 2009), an iterative procedure for estimating the model parameters (Dempster et al. 1977; Roweis 1997) that offers an easy way to handle incomplete datasets by estimating missing values alongside latent variables (Roweis 1997; Porta et al. 2005; Yu et al. 2010). This is particularly advantageous as most of the phylogenetic comparative methods handling missing values are restricted to low-dimensional datasets (e.g., (Clavel et al. 2015; Goolsby et al. 2017; Thorson and Van Der Bijl 2023) but see (Hassler et al. 2022)), or require the data to be imputed beforehand (e.g., for the phyPCA). Despite its iterative nature, because the EM primarily works on the reduced set of latent variables (lower-dimensional space), it often involves mathematical operations that are less computationally demanding than working directly on the observed traits. Other approaches based on continuous latent variable models for data dimension reduction have been adapted to the phylogenetic framework, such as the phylogenetic Factor Analysis (Tolkoff et al. 2018; Hassler et al. 2022), and a recently developed phylogenetic version of the PPCA (Caetano and Hearn 2026). Unlike the probabilistic PCA, the phylogenetic Factor Analysis assumes an anisotropic error (each latent variable can have its own noise contribution; see the Material and Methods section below), providing more flexibility to the model. However, this also renders it non-rotation-invariant, a condition required for analysing traits with no natural orientation in space such as Procrustes aligned geometric morphometric datasets (Rohlf 1999). Furthermore, the meaning of each factor (equivalent to the probabilistic principal components axes) can change as the number of selected factors changes (see (Tipping and Bishop 1999; Anderson 2003, 2004)). Finally, current implementations of the Phylogenetic Factor Analysis assume Brownian motion, although it could in theory be extended to other evolutionary models. Very recently, another phylogenetic version of the PPCA (Caetano and Hearn 2026) was independently proposed to extend the approach by (Tipping and Bishop 1999) for integrating the phylogenetic structure of various evolutionary model. This formulation allows to infer evolutionary parameters for different trait evolution models, and performs model selection to test evolutionary hypotheses (when *p<n*). However, this method cannot be applied to datasets containing missing values, which are commonplace in many large *omics* datasets, including 3D morphometrics involving fossils.

Here, we present a computationally efficient phylogenetic version of the probabilistic PCA (coined P3CA for Phylogenetic and Probabilistic PCA) to model the evolution of the latent variables along the branches of a phylogenetic tree. It scales to high-dimensional datasets and includes an EM algorithm that can handle missing data. We illustrate how this approach can simultaneously estimate the evolutionary parameters and the principal component axes by implementing the Pagel’s lambda model. This model enables the estimation of phylogenetic signal on high-dimensional traits through the parameter *λ* (Pagel 1999; Pearse et al. 2025). We use simulations to assess the accuracy in the parameter estimates obtained with the P3CA (i.e., the parameters of the continuous latent variables model, Pagel’s *λ* and missing values). Finally, we apply the P3CA to a high-dimensional 3D geometric morphometrics dataset (3669 coordinates) describing the skull shape of 24 extant and 19 extinct crocodyliforms species from Felice et al. (2021), with around 4% of missing values. We analyse the extent to which skull shape covaries with diet and habitat.

## Materials and Methods

### A Phylogenetic and Probabilistic PCA

Throughout the text, we use uppercase letters to refer to matrices and lowercase letters for scalars. *Tr*(***X***), |***X***|, ***X***^−1^, ***X***^*T*^, and ⟨***X***⟩, indicate the trace, determinant, inverse, transpose, and expected value of the matrix ***X***. We note the matrix dimensions by subscripts when these are not directly specified.

In the continuous latent variable model of the P3CA, the *n* × *p* matrix ***Y***, that describes the *p* traits for *n* species, is related to a lower *n* × *q* space ***Z*** of independent and unobserved variables through the matrix ***W***_p×q_ (Tipping and Bishop 1999; Li et al. 2009):

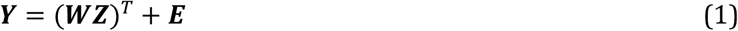

The matrix ***E***_n×p_ corresponds to the model error that captures the traits variance not included in the reduced space. We assume that ***Y, Z***, and ***E*** follow a matrix-variate normal distribution of the form ℳ𝒩_*a,b*_ (Θ_*a*×*b*_, **U**_*a*×*a*_, **V**_*b*×*b*_) with zero-mean (Θ = 0). Note that this formulation is equivalent to the multivariate normal distribution 𝒩_*ab*_ (*vec*(Θ_*ab*_), **V**_*b*×*b*_ ⊗ **U**_*a*×*a*_) where *vec*(Θ) is the vectorised form of the mean Θ and ⊗ describes the Kronecker product. For an easier explanation of the model, we will assume that ***Y*** is already centred. We therefore omit the mean in our formulation (but see below for its treatment).

Unlike the original PPCA (Tipping and Bishop 1999), in our phylogenetic PPCA framework, the *n* rows of ***Y*** are not assumed to be independent; rather, they may be correlated due to the shared evolutionary history of the species. We follow (Li et al. 2009) and assume that both the latent variables and the error term are phylogenetically correlated:

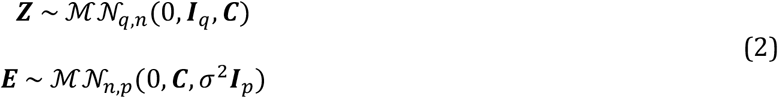

Where ***I*** is the identity matrix, and ***C*** the phylogenetic variance-covariance matrix corresponding to a given evolutionary model (such as Brownian-motion, the Pagel’s lambda model, or other models (Revell and Harmon 2008)). The parameter *σ*^2^ controls the relative proportion of noise contributing to the variance in the traits. The number of latent variables *q* is assumed to be known.

Conditional on the latent variable, the traits ***Y*** are also distributed according to a matrix variate normal distribution:

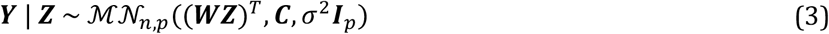

Integrating over the latent variable, the marginal distribution of ***Y*** is known in closed form (see (Li et al. 2009)):

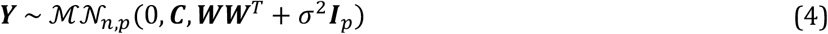

The marginal log-likelihood of ***Y*** is:

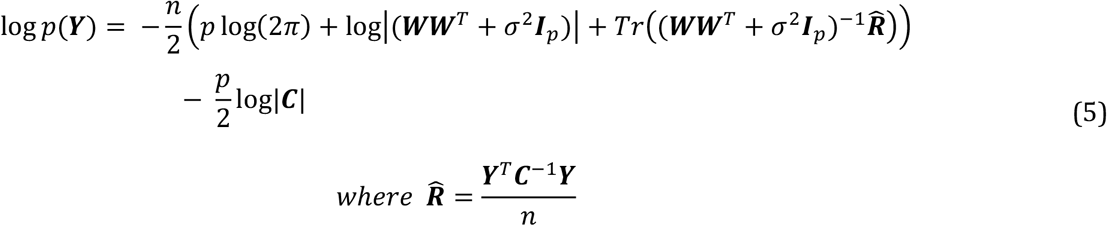

The maximum likelihood estimators for ***W*** and *σ*^2^, are given by Li et al. (2009):

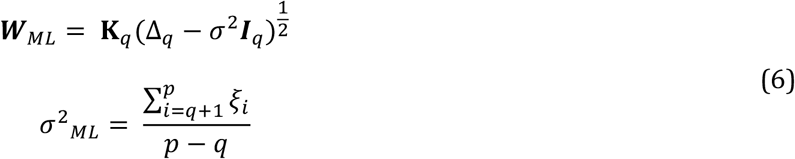

Where Δ is a diagonal matrix with the first *q* largest eigenvalues (*ξ*) of 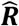, ***K*** is a *p* × *q* matrix containing the corresponding eigenvectors, and ***I***_*q*_ is a *q*×*q* identity matrix. As in the conventional probabilistic PCA, *σ*^2^ corresponds to the variance of all the discarded dimensions (*p* − *q*) (Tipping and Bishop 1999). Hence, when *σ*^2^ = 0 the P3CA converges to a phylogenetic PCA, although in practice the variance-covariance matrix ***WW***^*T*^ + *σ*^2^***I***_*p*_ would become singular when *p* > *n*, and the log-likelihood in equation (5) could not be evaluated. In addition, when ***C*** = ***I***_*n*_ the model in equation (1) converges to the conventional probabilistic PCA.

### Expectation – Maximisation (EM) algorithm

By maximising the marginal log-likelihood in equation (5) with the analytical solutions for ***W*** and *σ*^2^ (equation (6)) we showed that the evolutionary parameters can be correctly inferred (Figure S1, S2). However, this requires complete datasets. Therefore, we developed an alternative Expectation-Maximisation (EM) algorithm that can handle missing values.

The EM algorithm (EM hereafter) is an optimisation technique for parameter inference when the model includes latent or unobserved variables (Dempster et al. 1977). The EM aims to find the parameters estimates by maximising the marginal log-likelihood of the observed data (log *p*(***Y***), equation 5). Instead of optimising this likelihood directly, which can be computationally challenging, it operates by taking the expectation of the joint log-likelihood of the data (both observed ***Y***_***o***_ and missing ***Y***_***m***_) and the latent variables (log *p*(***Y, Z***), or log *p*(***Y***_***o***_, ***Y***_***m***_, ***Z***) when there are missing values, hereafter referred to as the complete-data log-likelihood). This expectation is computed with respect to the distribution of the latent variables (and with respect to the distribution of the missing values if any) conditional on the observed data and the current parameters (Dempster et al. 1977; Roweis 1997; Porta et al. 2005). The key idea underlying the EM is that working with the joint distribution, factorised as *p*(***Y, Z***) = *p*(***Y***|***Z***)*p*(***Z***), is often computationally more tractable than evaluating *p*(***Y***) directly. Essentially, it approaches the problem by breaking it down into two iterative steps: an expectation step (E-step) and a maximisation step (M-step). In the *E-step*, the expectation of the complete-data log-likelihood function (equation 9) is computed with respect to the conditional distribution of the latent variables (and missing data) given the observed data and current parameter estimates for ***W*** and *σ*^2^. Then, in the *M-step*, the parameter estimates maximising this expected complete data log-likelihood are estimated. The algorithm needs to be initiated with a set of values for ***W*** and *σ*^2^, and then iterates between the two steps until convergence (Dempster et al. 1977; Roweis 1997).

In our EM algorithm, we consider both the latent variables ***Z*** and the missing values in ***Y***, if any, as the unobserved variables to be estimated. The conditional distribution of the latent variable required for the *E*-step is given by (Li et al. 2009):

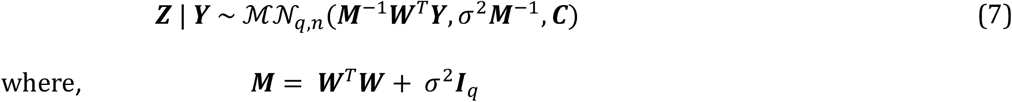

Because in comparative data the missing values are usually scattered throughout the matrix ***Y***, we treat the missing values for each trait separately, defining ***Y*** = [*y*(*x*), …, *y*(*x*_*p*_)] with *x* ∈ {1, …, *p*}, and where *y*(*x*) = (*y*_1_(*x*), …, *y*_*n*_(*x*))^*T*^ are the values of the trait *x* for the *n* species. We treat the missing values (*y*(*x*)_*m*_) as random variables normally distributed conditional on the observed values (*y*(*x*)_*o*_), the latent variables ***Z***, the parameters ***W*** and *σ*^2^.

The conditional distribution of the missing values for the trait *x* is given by:

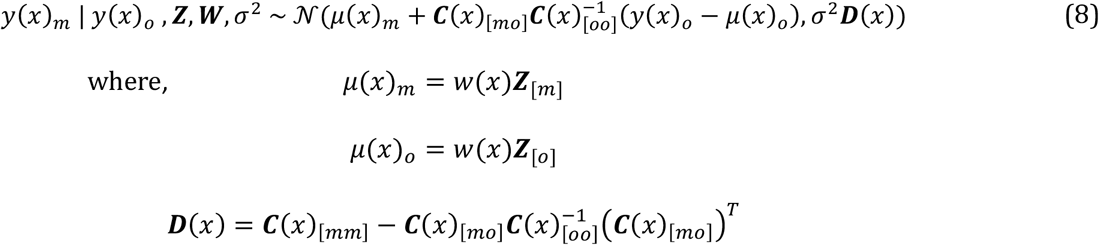

In equation (8), the subscripts *m* and *o* stands for the missing and observed values respectively, and *μ*(*x*), ***C***(*x*), and *w*(*x*), are the mean, the phylogenetic variance-covariance matrix, and the vector mapping the *q* latent variables to the trait *x*. An extended presentation and the complete derivation of the EM algorithm is described in Appendix I; below we summarize the two steps of the algorithm.

#### E-Step

The expectation of the complete-data log-likelihood with respect to the conditional distribution of the latent variables (*q*(***Z***)) and the missing values (*q*(***Y***_*m*_)) is given by:

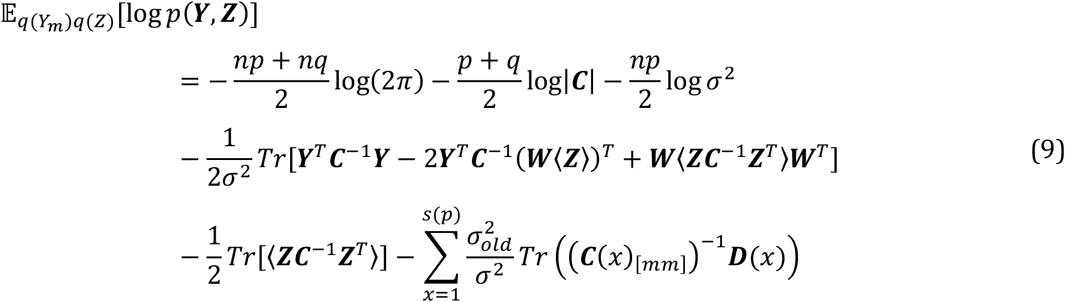

Where,

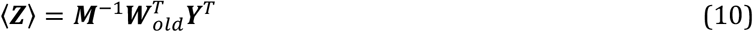

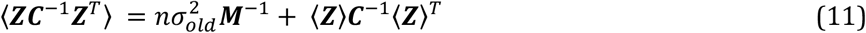

The subscript *old* indicates the value is from the previous iteration of the algorithm, and *s(p)* indicates the set of the *p* traits with missing values. In equation (9), the missing values in ***Y*** were filled-in by their expectation for each trait *x* (⟨*y*(*x*)_*m*_⟩), given by:

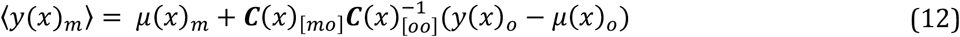

#### M-step

Maximising equation (9), we obtain the maximum-likelihood estimators for ***W*** and *σ*^2^ as:

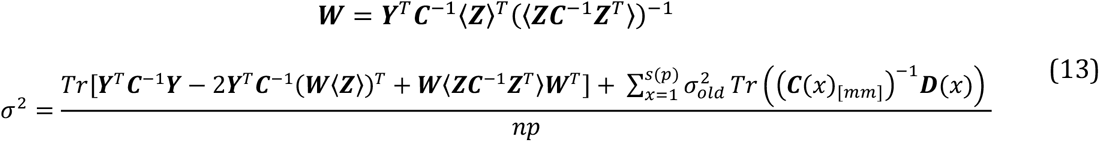

We assess the convergence of the EM by evaluating the marginal log-likelihood (equation 5) that we approximate using its lower bound (decomposed as the expected joint log-likelihood plus the differential entropy for the latent variables (*q*(***Z***)) and the missing values (*q*(***Y***_*m*_); see Appendix I, (Roweis and Ghahramani 1999)):

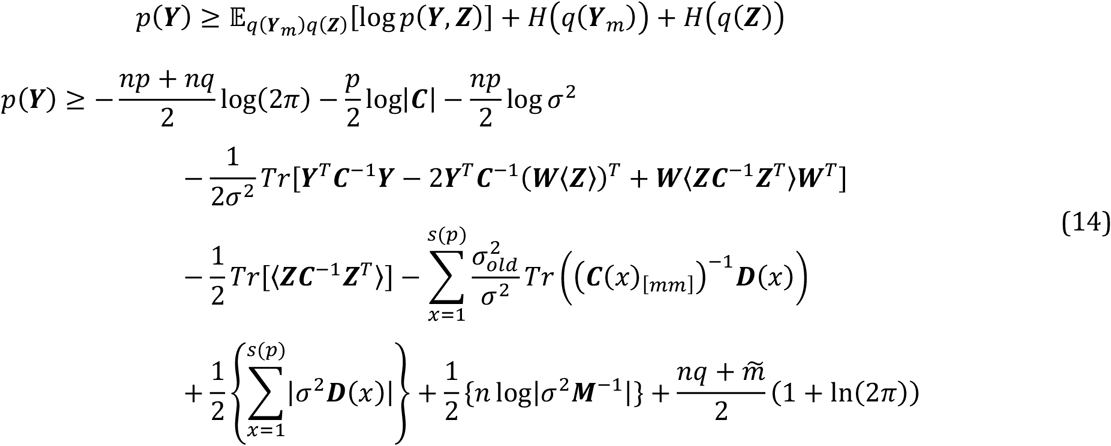

Where, 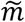 is the total number of missing values in ***Y***. Note that ***C*** is fixed inside the EM steps. We estimate the evolutionary parameters describing ***C*** (e.g., *λ* from Pagel’s lambda model), by maximising the lower-bound of the marginal log-likelihood of the data (i.e., equation 14 at convergence of the EM) (Appendix II). In the supplementary material we compare the estimators given by equation (13) at convergence to the analytical solutions of equation (6) when there are no missing values.

### Implementation

We implemented the P3CA with Pagel’s lambda model (Pagel 1999) in the *p3ca()* function of the R package ‘mvMORPH’ (Clavel et al. 2015). Pagel’s lambda model is convenient because it allows to describe both Brownian motion model (*λ* = 1) and deviations from it (*λ* ≠ 1). The estimation of Pagel’s lambda in the P3CA is done through the transformation of the matrix ***C***. For complete datasets, we optimise the *λ* parameter by maximising the marginal log-likelihood *p*(***Y***) (equation 5), and obtaining the maximum likelihood estimates for ***W*** and *σ*^2^, conditional on the transformed ***C*** matrix (***C***^*λ*^), with analytical solutions (equation 6). We provide an efficient algorithm to compute the marginal log-likelihood for complete datasets in Appendix (III). When the dataset includes missing values, we optimise *λ* (i.e., ***C***^*λ*^) by maximising the lower-bound of *p*(***Y***) (equation 14) with ***W*** and *σ*^2^ estimated by the EM (equation 13) (see Appendix II). In both cases, we use Brent’s algorithm (Brent 1971) implemented in the *optim()* function in R for the numerical optimisation.

The model in equation (1) assumes that the matrix ***Y*** is centred. In the P3CA, this centring is achieved by removing from ***Y*** the mean **Θ**, whose maximum-likelihood estimate (the ancestral state of each trait) is given by:

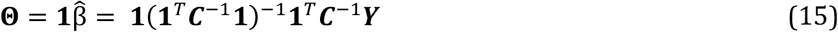

Where **1** is a one-column matrix (*n* × 1) mapping the ancestral states estimates 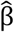 to each species. When there are missing values, the mean is calculated trait-wise using only the observed data, using a pruning algorithm (Felsenstein 1973). Because with small sample size (*n*) the parameter estimates (such as *σ*^2^ and *λ*) can be biased, we also implement a restricted maximum likelihood (REML) version. This is done by replacing the number of species *n* by *n* − 1 for all the estimators and by adding the term |**1**^*T*^***C***^−1^**1**|^*p*^ to the likelihood function (Patterson and Thompson 1971). The REML for the lower-bound of the marginal log-likelihood of the data *p*(***Y***) is given by:

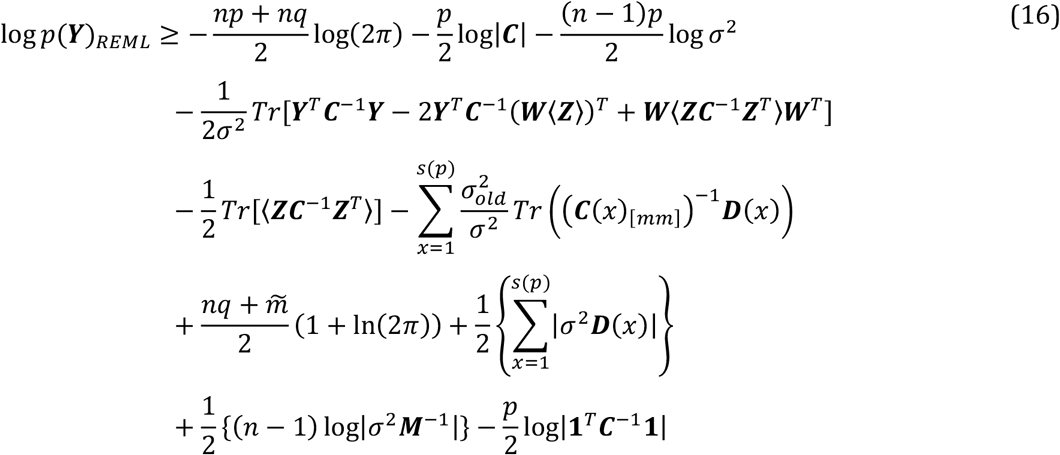

Note that, as in conventional probabilistic PCA, *q* is fixed *a priori* during the optimisation of all parameters. Approaches for selecting the number of latent variables *q* are discussed in the *Discussion* section below.

### Assessing the performances of the P3CA

We evaluated the performances of the P3CA with trait datasets simulated according to the continuous latent variable model described in equation (1). First, we built the matrix ***W*** as ***W*** = ***U***(**Λ**^0.5^)***R***, where ***U*** and ***R*** are *p* × *q* and *q* × *q* matrices of eigenvectors respectively, sampled from a uniform distribution on a Stiefel manifold, thereby ensuring the required orthonormality (*rustiefel()* function in ‘rstiefel’ package in R (Hoff and Franks 2021)). **Λ** is a diagonal matrix with *q* eigenvalues sampled from a log-normal distribution with 0-mean and 1-standard deviation, generating heterogeneity in the variance explained by the latent variables *(rlnorm()* function from base R). We scaled the eigenvalues so that approximately 80% of the variance in the dataset is contained in the *q* latent variables. We simulated the phylogenetic variance-covariance matrix ***C***, using Pagel’s lambda model for a range of *λ* value. To achieve this, we first simulated a phylogenetic tree under a pure-birth process (*pbtree()* function in ‘phytools’ package in R (Revell 2012, 2024)) and then transformed it according to the *λ* value (*rescale()* function in ‘phytools’). Using the transformed tree, we simulated the latent variables ***Z*** as *q* independent Brownian traits with 0-mean and unit variance using the *mvSIM()* function in ‘mvMORPH’. Similarly, the error term (***E***) was simulated as *p* independent Brownian traits with 0-mean and variance *σ*^2^ on the transformed tree in the function *mvSIM()*. In all our simulations, we used 50 species (*n*), 5 latent variables (*q*), and either 25 or 100 variables (*p*), corresponding to the low-dimension (*n* > *p*) and high-dimension (*n* < *p*) cases, respectively. We simulated datasets under *λ* = 0.98, which correspond to traits evolving almost as Brownian motion (BM), and *λ* = 0.6, which represent a substantial deviation to BM. We set *σ*^2^ = 0.1, which represent about 20% of the total variance in the dataset (i.e., the variance not explained by the *q* latent variables). For each combination of parameters, we randomly removed 2.5% and 5% of the observations to generate datasets with missing cases. We simulated 100 datasets for each set of parameters (*p* = low/high-dimension, *λ* = 0.6/0.98, *missing* = 0%, 2.5%, 5%). We then used these simulated data for assessing the accuracy of the approach in terms of i) parameter inference; ii) lower-dimensional space reconstruction; and iii) missing values estimation. For (ii) and (iii), we also compared the performance of the P3CA to other PCA approaches.

#### i) Parameter inference

We fitted the P3CA with REML to each dataset using the EM algorithm and compared the difference between the inferred and simulated *λ* and *σ*^2^ parameters.

#### ii) Reconstruction of the lower-dimensional space

We evaluated the accuracy of the lower-dimensional space reconstruction by comparing the estimated and simulated matrices ***W***. For the comparison we calculated the largest principal angle between the subspaces (Bjorck and Golub 1973) spanned by both matrices using the *angle()* function in the ‘rospca’ package (Reynkens 2024). The standardised angle ranges from 0 to 1. The closer the value is to 0, the closer the subspaces are (i.e., subspaces are equal when the angle is 0). We also evaluated the lower-dimensional space through the correlation analysis between the *q* estimated principal components to the simulated ones.

We compared the lower-dimensional space obtained using the P3CA with those obtained using conventional PCA and phylogenetic PCA (phyPCA), two of the most widely used approaches in studies of phenotypic evolution. To do this, we performed a conventional PCA using centred traits in the *prcomp()* function from the base R package, and a phyPCA using the *phyl*.*pca()* function in the ‘phytools’ package in R. We used the phyPCA with Pagel’s lambda model for the low-dimensional datasets (*n* > *p*), and with the Brownian-motion model for the high-dimensional datasets (*n* < *p*), because it is the only model directly applicable in this case. Then, we calculated the last standardised principal angle between ***U***, the matrix of eigenvectors used to generate ***W*** in the simulations, and the matrix of eigenvectors obtained by each approach. We also compared the approaches by estimating the correlation between their *q* first principal components and the simulated ones (i.e., the centred traits ***Y*** rotated by simulated ***W***). As both the PCA and the phyPCA do not allow for missing values, this comparison was restricted to the complete datasets.

#### iii) Missing values estimation

Finally, to evaluate the accuracy of the estimation of the missing values, we calculated the mean squared error between the values that were removed from the datasets (*missing* = 2.5% / 5%) and the values estimated by the P3CA. We compared the P3CA to PhyloPars (Bruggeman et al. 2009), a phylogenetic data-imputation approach. We used PhyloPars under a Pagel’s lambda model, and using the restricted maximum-likelihood (REML) (*phylopars()* function in ‘Rphylopars’ package in R (Goolsby et al. 2017)). This comparison is restricted to low-dimensional datasets (*n* > *p*) since PhyloPars is based on maximum likelihood and cannot handle high-dimensional datasets.

### Application of the P3CA to empirical data: the skull shape morphospace of Crocodyliformes

We used the morphospace of skull shape in Crocodyliformes to illustrate the differences when (1) a Brownian-motion model – the only currently available model for phyPCA on high-dimensional datasets – is used, and when (2) deviation from this model is allowed using the P3CA with the Pagel’s lambda model. We did not make a comparison to the recently developed phylogenetic PPCA of (Caetano and Hearn 2026), because it is not possible to fit Pagel’s lambda model to a high-dimensional dataset such as the one used here with this implementation. Hereafter, we refer to morphospace as the reconstruction of the lower-dimensional space. We used the high-dimensional dataset published by (Felice et al. 2021) composed of 3996 coordinates (1332 landmarks and semi-landmarks in 3D) describing the right-side of the skull for both, extant (24) and extinct (19) lineages. This dataset contains around 4% of missing values, concentrated in five extinct lineages (*Anatosuchus minor, Caipirasuchus stenognathus, Cricosaurus* sp., *Pholidosaurus* sp., *Sarcosuchus imperator*). Because the extinct lineages provide evidence of contrasting skull shape morphologies throughout the phylogeny, the reconstructed morphospaces can show different patterns depending on the trait evolution model considered. This dataset can be analysed with the currently-available dimensionality reduction approaches (PCA and phyPCA) if the missing values are imputed beforehand. Since there is no phylogenetically informed data-imputation approaches directly applicable to high-dimensional datasets, we followed the common practice in the field, i.e. we used a non-phylogenetic imputation technique, as in the original study (thin-plate spline interpolation (Bookstein 1989), performed with the *fixLMtps()* function from the ‘Morpho’ package in R (Schlager 2017)). We used one of the phylogenetic trees from the original study of Felice et al. (2021) (see Supplementary material - *Exploring the morphospace for the skull shape in Crocodyliformes*); but it should be noted that the phylogenetic relationships within Crocodylomorpha are still debated with no consensus (e.g., see a complete review in (Rio and Mannion 2021; Darlim et al. 2022; Burke et al. 2024)).

To obtain the phyPCA morphospace under the Brownian-motion model (strategy 1), after imputing the missing values as described above, we performed a generalised Procrustes superimposition (*ProcGPA()* function from ‘Morpho’) on the filled dataset, and used the superimposed data to perform the phylogenetic PCA (*phyl*.*pca()* from ‘phytools’). To obtain a morphospace from the P3CA with Pagel’s lambda model (strategy 2), we first aligned the original dataset through a generalised Procrustes superimposition with missing values (*align*.*missing()* from the ‘LOST’ package in R (Arbour and Brown 2025)). The approach implemented in *align*.*missing()* performs the superimposition of the incomplete specimens to a consensus conformation built from the complete specimens (Arbour and Brown 2014). We fitted the P3CA with REML to the superimposed landmarks coordinates using the EM and considering the first four dimensions (*q* = 4). The EM was initiated with the analytical solution for ***W*** (equation 6) obtained on the dataset with missing values replaced by 0, and by setting *σ*^2^ = 0.5. We compared the morphospaces obtained using the two strategies by examining the patterns obtained across the principal components (PC) axes and the loading matrix. The loading matrix represents the correlations between the traits and the PC axes, enabling the identification of the main features contributing to each axis. To interpret the PC axes, we used the information on diet and habitat, two factors potentially influencing skull evolution (e.g., (Pierce et al. 2008; Ballell et al. 2019; Godoy 2020; Felice et al. 2021)), provided in the original study (Felice et al. 2021). These data categorise the species into five ecological groups: carnivores (32) and piscivores (2) in freshwaters, carnivores (1) and omnivores/herbivores (6) in terrestrial systems, and piscivores in marine environments (2). We assessed the extent to which lineages with similar ecologies cluster together in morphospace.

## Results

### P3CA performances

#### (i) Parameter inference

Using the P3CA we were able to infer *λ* with high accuracy regardless of its value, the number of variables (low/high dimension) and the percentage of missing values (up to 5% of missing values) (Figure 1a). *λ* estimates were most accurate close to value of 1 (Figure S1), and slightly overestimated when *σ*^2^ was close to 0 (Figure S2*a*). *σ*^2^ was slightly underestimated across all the evaluated conditions, regardless of the number of variables or the percentage of missing values (Figure 1b). The slight underestimation of *σ*^2^ increased for larger values of this parameter, but this did not affect *λ* estimates (Figure S2*b*).

**Figure 1.**
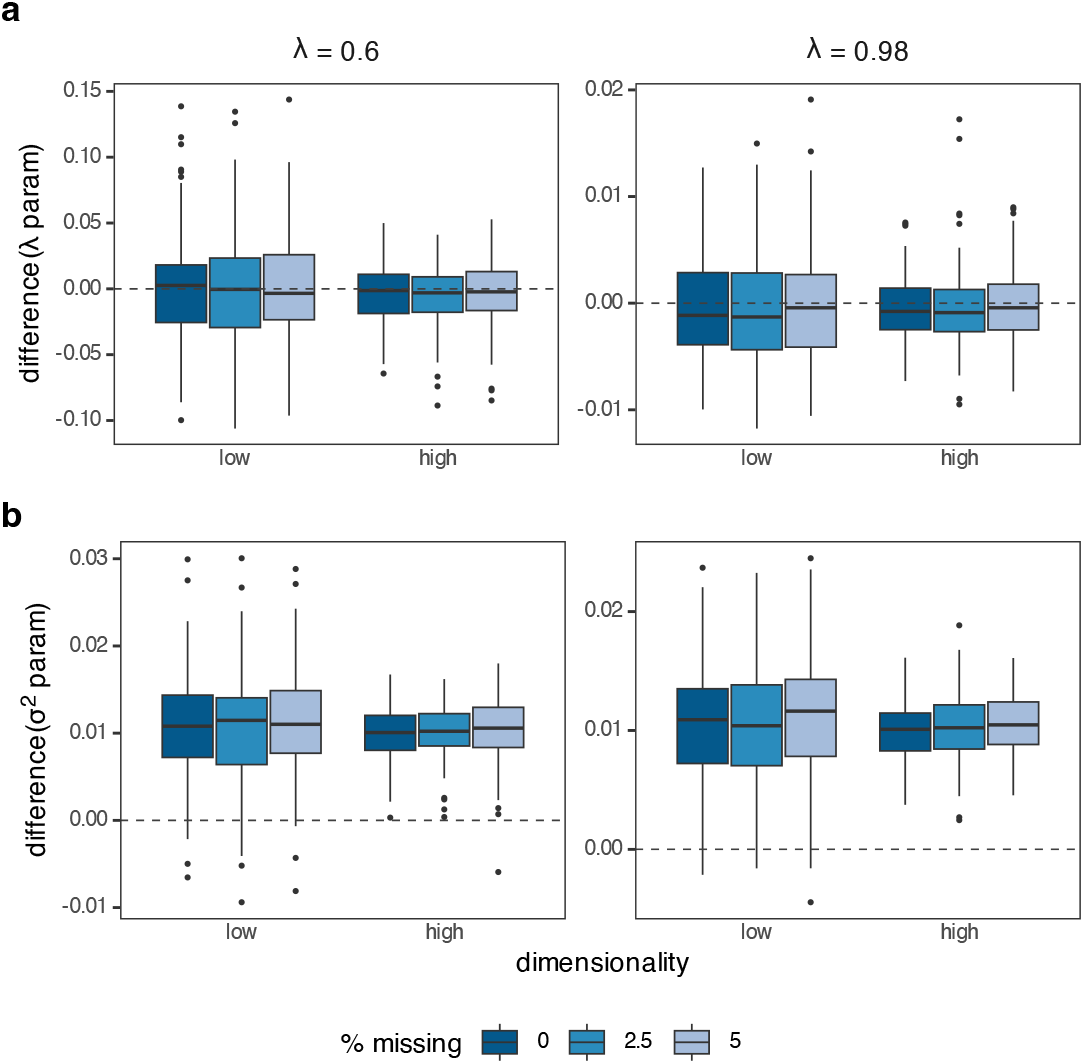
Difference between the simulated and inferred value for the parameters (**a**) *λ* and (**b**) *σ*^2^, for traits simulated under *λ* = 0.6 (left panels) and *λ* = 0.98 (right panels) (Brownian motion model when *λ* = 1). *λ* is inferred accurately and *σ*^2^ is slightly underestimated. The inference is consistent regardless of the percentage of missing values, the dimensionality (low/high-dimension) and the value of *λ* used in the simulations. The figure shows the inference (from 1^st^ to 3^rd^ quantile in boxes, while median in solid line) across 100 simulations with *n* = 50, *q* = 5, *σ*^2^ = 0.1, and *p* = 25 (low-dimension) or *p* = 100 (high-dimension).

#### (ii) Comparison of the lower-dimensional space reconstruction between PCA approaches

The P3CA performed the best at recovering the lower-dimensional space across simulated conditions. Moreover, the similarity between the matrix ***W*** estimated by the P3CA and the simulated one was independent of the number of variables, the percentage of missing values and the value of *λ* (Figure 2a). Similar results were obtained through the comparison of the correlation between the principal components, with the correlation decreasing from the first to the last component evaluated (Figure S4). Compared to other approaches, the P3CA always outperformed the conventional PCA in terms of ***W*** estimates (smaller angles between the simulated and estimated ***W*** matrices). The P3CA had similar performances to the phylogenetic PCA (phyPCA) when the latter could be fitted using Pagel’s lambda model (i.e., in low-dimensional situations), and performed better otherwise (Figure 2b, Figure S3). Finally, the principal components obtained by the P3CA showed the highest overall correlation with the simulated ones. In low-dimensional datasets, the performance was comparable to the phyPCA fitted under Pagel’s lambda (Figure S5). In high-dimensional datasets, the performance was similar to the phyPCA fitter under BM only when *λ* value was close to 1 (Figure S6).

**Figure 2.**
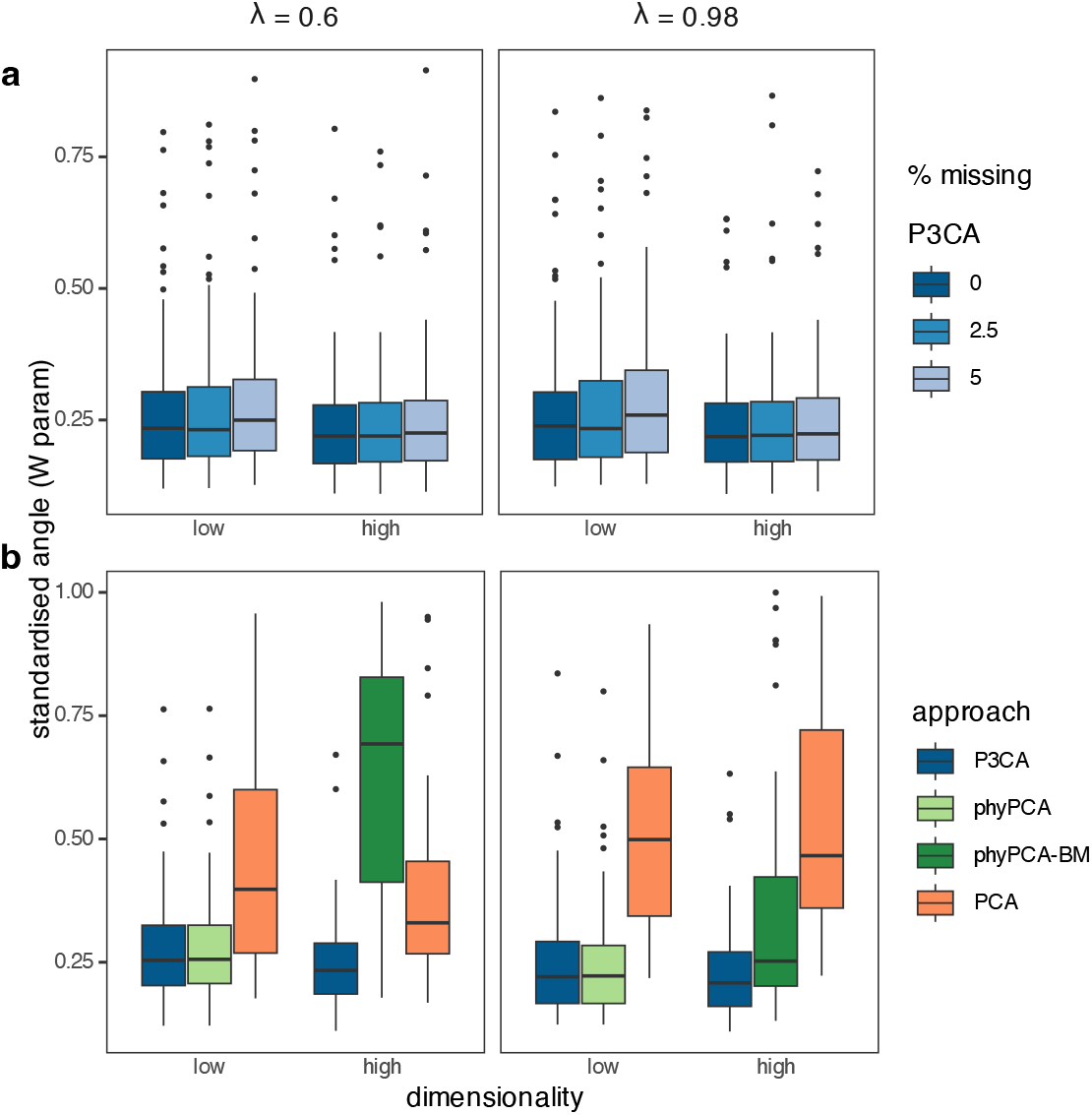
Standardised angle between the simulated and inferred factor matrix ***W***, for traits simulated under *λ* = 0.6 (left panels) and *λ* = 0.98 (right panels) (Brownian motion model when *λ* = 1). The closer the angle to 0, the closer the matrices. (**a**) When ***W*** is inferred from the P3CA, the angle is similar across datasets simulated under different *λ* values, in low and high dimension, and with varying percentages of missing values. (**b**) When compared to other approaches, the angle is the smallest for ***W*** inferred from the P3CA. In low-dimension, the phylogenetic PCA (phyPCA) can be fitted under Pagel’s lambda model, and it then performs as well as the P3CA; in high-dimension, the phyPCA can only be fitted under the Brownian-motion model (phyPCA-BM), and the P3CA outperforms all other approaches. The figure shows the inference (from 1^st^ to 3^rd^ quantile in boxes, while median in solid line) across 100 simulations with *n* = 50, *q* = 5, *σ*^2^ = 0.1, and *p* = 25 (low-dimension) and *p* = 100 (high-dimension).

#### (iii) Estimation of the missing values

Across all simulated conditions, the P3CA produced a smaller mean squared error (MSE) between the estimated and the simulated missing values than the maximum-likelihood implementation in RphyloPars (Figure 3). For both the P3CA and PhyloPars, smaller MSE were obtained with higher values of *λ*.

**Figure 3.**
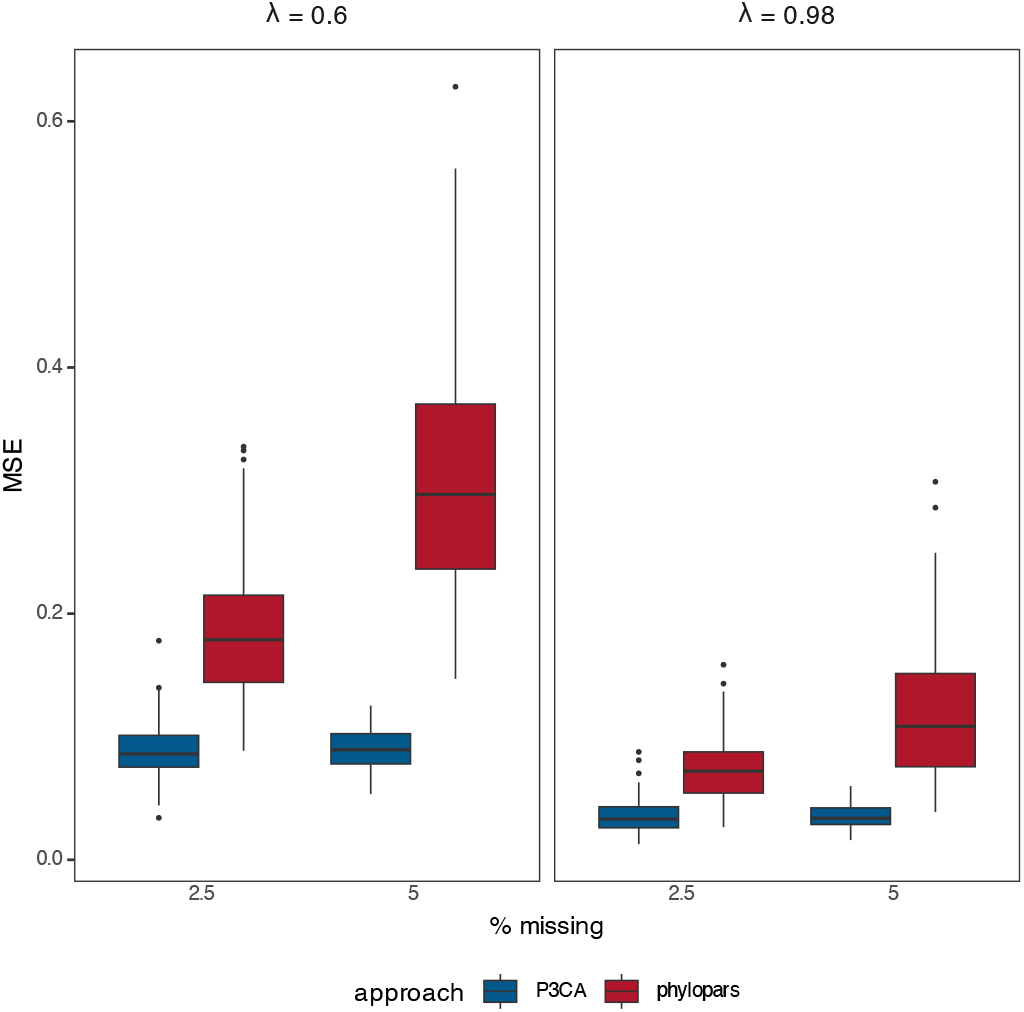
Mean square error (MSE) between simulated and estimated missing values, for traits simulated under *λ* = 0.6 (left panels) and *λ* = 0.98 (right panels) (Brownian motion model when *λ* = 1), and estimates obtained with the P3CA and PhyloPars, in datasets with 2.5% and 5% incompleteness. The error is lower for the P3CA across the two values of *λ* used in the simulations. The figure shows the inference (from 1^st^ to 3^rd^ quantile in boxes, while median in solid line) across 100 simulations with *n* = 50, *q* = 5, *σ*^2^ = 0.1, and *p* = 25 (low-dimension).

### The skull shape morphospace of Crocodyliformes

Using four latent variables, around 64% of the variance in the skull shape is explained and the estimated *λ* is 0.81. Despite a rather high *λ* (Brownian-motion model is *λ* = 1), the P3CA and the phylogenetic PCA under Brownian motion (phyPCA-BM) substantially differ in their morphospaces. Only the first component is congruent, while the other axes differ, changing the possible ecological interpretation in terms of associations with diet and habitat regimes (Figure 4a). We describe here the first and second component, and the third and fourth are described in the Supplementary material (see *Exploring the morphospace for the skull shape in Crocodyliformes*). With both methods, the first component describes a gradient ranging from shortened, wide and tall skulls with an expanded pterygoid, to long, narrow and flattened skulls with a smaller, more contracted pterygoid (Figure 4b, Figure S7). This matches a gradient from omnivorous/herbivorous to piscivorous lineages (left to right in Figure 4a). The second component describes important changes in the eye position (frontal and prefrontal area) and the occipital condyle with the two methods, but differences appear in the contribution of other anatomical areas. For example, the P3CA detects variations around the pterygoid and the relative position of the quadrate to the occipital condyle (i.e., varying from skulls with pterygoids more vertically oriented and quadrates farer to the occipital condyle, to skulls with pterygoids facing backwards and quadrates closer to the occipital condyle), while the phyPCA-BM detects variations related to the shape of the snout (Figure 4c). The second component reinforces the separation between omnivorous/herbivorous and piscivorous lineages with the phyPCA-BM, but not with the P3CA. Instead, the P3CA clusters terrestrial and marine lineages at one extreme, with no clear separation of freshwater lineages which occupy a wider space (Figure 4a).

**Figure 4.**
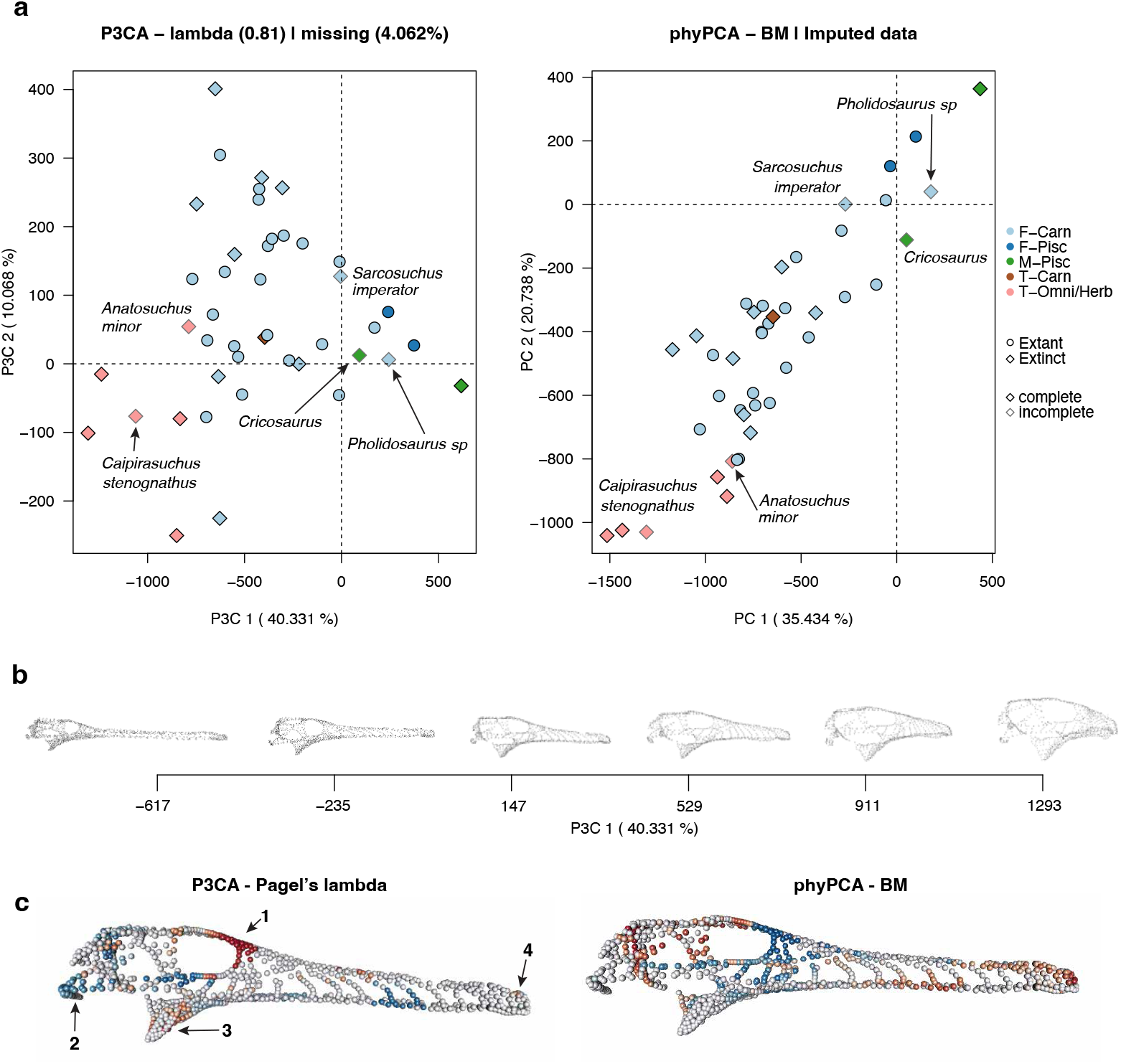
(**a**) Comparison of the morphospace obtained for skull shape in Crocodilians with the P3CA under Pagel’s lambda model (left panel), and with the phyPCA under the Brownian-motion model (right panel). For the phyPCA -BM, a non-phylogenetic data-imputation was performed beforehand. While the first component captures similar shape variation across the two methods, the second one differs markedly. The strong segregation aligned to the habitat regime shown in the second component of the phyPCA -BM is not evident in the second component of the P3CA; terrestrial lineages cluster toward one extreme, but freshwater lineages are broadly distributed, with no clear separation between the two habitats. (**b**) Variation in skull shape along the first component obtained with the P3CA. The figure shows the projection on the first axis of landmarks and semi-landmarks into a mean shape(**c**) Traits (landmarks and semi-landmarks) with the highest variation along the second component under the P3CA (left panel) and the phyPCA -BM (right panel). The coloured landmarks/semi-landmarks are those with load values greater than 0.2 or less than -0.2. The threshold (0.2/-0.2) is defined purely for visualisation purposes. The loads, from the loading matrix, correlate the component (latent variable) with each trait (landmark or semi-landmark). The stronger the load, the greater the contribution of the trait to the PC axis. The frontal and pre-frontal areas (1) and the region around the occipital condyle (non-visible) are important contributors of variation along the second axis under both methods. The quadrate (2) and pterygoid (3) appear as important contributors only with the P3CA, and in contrast, changes around the snout (4) are only detected with the phyPCA -BM. Panel (c) shows the projection of the landmarks and semi-landmarks onto a common mean shape.

## Discussion

We developed the probabilistic and phylogenetic PCA (P3CA), a latent variable based dimensionality reduction technique for modelling trait evolution on multivariate, including high-dimensional, datasets. The simulations support an accurate inference of the phylogenetic signal (Pagel’s *λ* parameter) for both low and high-dimensional datasets, even when those include up to 5% of missing values. This holds true irrespective of the slight underestimation of the parameter *σ*^2^, describing the trait variation not captured by the latent traits. Furthermore, the reduced space obtained with the P3CA is more accurate than that obtained using other dimensional reduction techniques, such as the conventional and the phylogenetic PCA. Finally, the P3CA is more accurate at estimating missing values than some current phylogenetic data imputation approaches, with the advantage of being applicable to high-dimensional datasets. Using the P3CA, we were able to fit a multivariate Pagel’s lambda model to an incomplete (around 4% of missing values) high-dimensional dataset describing the shape of the skull of living and extinct crocodyliforms.

### Degining the number of latent variables or PC axes

A common challenge when applying dimensionality reduction techniques is defining the number of dimensions *q* (PC or latent variables) for analysing the data. The P3CA requires the selection of *q* latent variables *a priori*. However, compared to other approaches such as the Factor Analysis, the lower-dimensional space reconstructed by the probabilistic PCA is not dependent on the number of latent variables used to fit the model – either using the analytical solutions or the EM. From a biological perspective, this implies that interpreting each latent variables or the reduced space is not affected by the choice made *a priori* for the number of latent variables. Only *σ*^2^ will vary, as expected, with the number of retained latent variables. The P3CA shares this property, assuming that each latent variable has evolved under the same process. However, *λ* estimation may vary depending on the number of latent variables used and whether the processes governing these variables differ (see the discussions on models that could relax the assumption of homogeneous processes below). In this case, an average phylogenetic signal will be estimated, which may impact the reduced space obtained.

Although we did not explore this aspect in our study, multiple approaches for selecting the number of relevant PC axes in the classical PCA or PPCA are also suitable for the P3CA. For example, one can use *ad-hoc* rules based on the eigenvalue distribution such as the Cattell’s scree-test or the Hull method (Cattell 1966; Jolliffe 2002; Ahn and Horenstein 2013), techniques based on information criteria such as those proposed by (Bai and Ng 2002, 2019), or resampling-based methods such as bootstrapping or cross-validation (e.g., (Krzanowski 1987; Josse and Husson 2012; Owen and Wang 2016; Dobriban and Owen 2019)). The performances of these techniques often vary with the data structure, as shown by study in multiple fields (e.g., (Deng and Craiu 2023)). In addition, some approaches have attempted to select the number of latent variables jointly during model fitting, and these could potentially be explored and implemented in the P3CA. For instance, the Bayesian PCA (Bishop 1998), and approaches using Bayesian variable selection (e.g., (Minka 2000; Nyamundanda et al. 2010; Bouveyron et al. 2018)) or penalisation methods (e.g., (Deng and Craiu 2023)), which jointly estimate the parameters describing the model along with the number of latent variables, are promising directions to investigate. However, these approaches often incur a high computational cost due to the complexity in solving these models (e.g., the integrals in these Bayesian formulations; (Bishop 1998; Minka 2000; Nyamundanda et al. 2010), or due to the need to perform the maximisation multiple times (Bouveyron et al. 2018; Deng and Craiu 2023). This is a research direction that we intend to pursue in the future.

### Parameter inference

The estimation of *λ* by the P3CA was accurate and unbiased while the parameter *σ*^2^ was slightly underestimated. The bias in *σ*^2^ however, is expected given that its estimation relies on the eigenvalues of the discarded dimensions (i.e., the *p* − *q* dimensions not considered in the lower-dimensional space). A general phenomenon when using the maximum-likelihood estimate of the trait covariance matrix (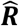, equation 5), is the degeneration of its eigenvalue structure as *p* approaches *n*. In these conditions, the largest eigenvalues of 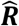 increase and the smallest one decreases (see for example, Passemier et al. 2017; Ledoit and Wolf 2020). Since *σ*^2^ is based on the discarded dimensions, which all include the smallest eigenvalues, underestimated eigenvalues directly result in an underestimation of *σ*^2^. With a large *n*/*p* ratio, this bias is expected to vanish. Indeed, in our simulations in low-dimension (*n* > *p*), the accuracy in the estimation of *σ*^2^ increases with larger *n* (Figure S8). These biases constitutes a known challenge for many phylogenetic comparative approaches that attempt to correct the distribution of eigenvalues of the covariance matrices (see (Clavel et al. 2019; Montoya et al. 2026)). Similarly, the resulting overestimation of the variance explained by the reduced space (Figure S9) also affects most approaches that rely on the eigenvalue structure of the sample covariance matrix. These include the P3CA, the probabilistic PCA, conventional PCA and phylogenetic PCA (e.g., Johnstone and Lu 2009; Passemier et al. 2017; Johnstone and Paul 2018). Different approaches have been proposed to alleviate the problem in the probabilistic PCA, by either constraining parameter estimates (e.g., Lopes and West 2004; Lee et al. 2012; Ma 2013) or directly correcting *σ*^2^ (Passemier et al. 2017). These approaches could probably be adapted for use with the P3CA and this should be considered in the future.

The bias in *σ*^2^ does not appear to affect the inference of the evolutionary parameter (*λ*), however *λ* estimates are less accurate when *σ*^2^ approaches zero. This is probably due to the instability of the covariance matrix describing the continuous latent model (***WW***^*T*^ + *σ*^2^***I***_*p*_), equations 4 and 5) when *σ*^2^ tends towards zero. The larger the difference between the largest and smallest eigenvalues of the matrix, the stronger the matrix instability (i.e. high condition-number), which is known to affect parameter estimation (e.g., (Jhwueng and O’Meara 2020)). The smaller *σ*^2^, the greater the difference in the eigenvalues of ***WW***^*T*^ + *σ*^2^***I***_*p*_, and thus the variance in the data modelled by the *q* latent variables (largest eigenvalues), results in an unstable parameter estimation and often an overestimation of *λ* (see for instance ML inference when *p*/*n* is close to 1 in Fig. S5 of (Clavel et al. 2019)).

### Performance of the EM algorithm

Our EM algorithm performed well, converging to the analytical solution for both *σ*^2^ And ***W***. Because neither of the E- or M-steps involve computations scaling with the large *p* × *p* covariance matrix usually used in phylogenetic comparative methods, the EM can theoretically scale to very high-dimensional datasets (with more than a thousand traits) when using a few numbers of latent variables *q*. However, as with any EM algorithm, two aspects require a particular attention. First, although the EM provides a stable convergence increasing monotonically in the likelihood surface, the procedure becomes slower as the number of parameters increases (Dempster et al. 1977; Wu 1983; Gentle 2009; Kuroda and Geng 2023) because the required number of iterations usually increases with the proportion of missing values and dimensionality (*p* and *q*). Second, since EM algorithms can converge to local rather than global maxima (Wu 1983; Tipping and Bishop 1999; Gentle 2009; Wu et al. 2017), the choice of initial values is crucial. In our simulation analyses, the EM converged to the analytical solution by starting from random values in both low and high-dimensional datasets. For the crocodyliform dataset however, the algorithm was faster and consistently converged at higher log-likelihood values when initiated with a well-chosen ***W*** matrix rather than with random ones (results not shown). For instance, analytical solutions for ***W*** on datasets where missing values have been filled by 0 or the arithmetical mean may be used. Although this strategy increases the processing time due to the eigen-decomposition it implies, it ensures stable convergence.

A major advantage of the EM algorithm is to handle datasets with missing values. It allows parameter inference even when datasets are incomplete (Roweis 1997), and as a by-product provides accurate estimates of the missing values (e.g., (Musil et al. 2002; Yu et al. 2010; Josse et al. 2011; Malan et al. 2020)). Our results show that the missing value reconstructions were better with the P3CA than with another phylogenetic imputation approach, in which the model parameters are estimated using ML alongside the data imputed by their conditional expectations. The difference in performance between both approaches is probably related to the number of parameters estimated that scale with dimensionality (*q* for P3CA and *p* for RphyloPars). In our simulations, an increase in the proportion of missing values, when distributed uniformly at random, had no effect on the accuracy of the inferred *λ* parameter and missing values. However, we can likely expect a loss of accuracy when this assumption is not met (e.g., when the missing values are clustered in some traits or clades). Some approaches have recently been proposed for the conventional probabilistic PCA to address the particular case of non-random missing values (Sportisse et al. 2020), but further work is needed to adapt these approaches to our implementation and assess their performance.

### Extending the tools for studying phenotypic evolution

A clear advantage of the P3CA compared to other phylogenetic PCAs (e.g., Revell 2009; Caetano and Hearn 2026) is that it allows to directly analyse incomplete high-dimensional dataset without filling missing values beforehand. Indeed, the current practice with high-dimensional datasets with missing values is to perform imputation using non-phylogenetically informed approaches before analysing these imputed datasets with PCM approaches (e.g., (Clavel et al. 2019; Goswami et al. 2022)). However, this pipeline might be prohibitive for very large datasets. Furthermore, the effect of using non-phylogenetic imputation (which are known to be less precise than phylogenetic ones, e.g., (Penone et al. 2014; Jardim et al. 2021; Johnson et al. 2021)) in model selection remains unclear.

The P3CA and other phylogenetic PCAs (Revell 2009; Caetano and Hearn 2026) are otherwise similar in that they both seek the axes with the largest variance in a multivariate trait evolutionary process (i.e., the main evolutionary changes) rather than in the raw trait variance. This implies that the P3CA axes are not orthogonal (i.e., they can be correlated); they still contain phylogenetic signal, and must therefore be analysed using phylogenetic comparative methods when they are used in subsequent analyses (Revell 2009; Uyeda et al. 2015; Clavel and Morlon 2020). A complete description of the statistical properties of the phylogenetic PCA can be found in (Revell 2009) and (Polly et al. 2013).

For high-dimensional data, our approach offers the advantage of allowing deviations from BM over the phylogenetic PCA (phyPCA). As we have shown using the crocodyliform skull shape, the morphospace can differ significantly depending on the fitted evolutionary model, even when the parameter *λ* is close to 1 (and therefore to BM). In our comparison, we found a general elongation and dorsoventral compression of the skull, associated mainly to the dietary regime differences, as well as changes in the eye position related to the habitat for both models (BM and Pagel’s lambda). However, while the second component reinforces this association under a BM model, removing the BM constraint by using Pagel’s lambda attenuates the strong link between morphology and diet. Instead, habitat regimes emerge as another potentially important ecological factor associated with the main morphological changes. Some of the features changing along the second component under Pagel’s lambda, such as the position of the quadrate relative to the occipital condyle and pterygoid, have been related to increased jaw movements complexity typical to a terrestrial life-style (as illustrated in the association of bit force to muscle pull direction in (Busbey and Thomason 1995; Srinivas et al. 2025)). We do not attempt to explain the evolution of the Crocodyliform skull shape given the small sample size and the ongoing debate on the phylogenetic relationships. Rather, we use this to illustrate how constraining the dimensionality reduction to the BM could lead to the overlooking of potentially relevant ecological factors. Furthermore, different results and potentially different conclusions may be reached when the PC axes are employed in subsequent statistical analyses, given that the changes explained by each PC axis differ between the evolutionary models.

Besides the aforementioned advantages, we believe that extending the phylogenetic dimension-reduction techniques will facilitate the study of the very large multivariate datasets generated in the last few decades. The computational complexity of the P3CA scales with lower dimensions (either *p* × *q* or *n* × *q*) than classical multivariate phylogenetic comparative methods; which have to deal with operations on *p* × *p* matrices. Furthermore, some computational and mathematical tricks can be applied to speed up inference even when using the ML solutions at large dimensions (equations 5 and 6; see Appendix III). Therefore, with the P3CA, it is theoretically possible to fit models to datasets containing tens of thousands of traits. To date, large datasets are typically reduced to the axes obtained from PCA or phyPCA-BM (geometric morphometric datasets comprising few thousand of traits, e.g., (Felice et al. 2019, 2021; Watanabe et al. 2019; Fabre et al. 2020, 2021; Coombs et al. 2022; Goswami et al. 2022)), or analysed under the assumption of trait independence (gene expression data comprising few tens of thousands, e.g., (Brawand et al. 2011; Chen et al. 2019; Flanagan et al. 2026)). However, these practices are known to lead to misleading results (see for instance (Revell 2009; Uyeda et al. 2015; Clavel and Morlon 2020)). Phylogenetic Factor Analysis (PFA), another phylogenetic dimensional reduction technique share with the P3CA the property of working from a reduced space and can be used on high-dimensional datasets (Tolkoff et al. 2018; Hassler et al. 2022). However, as the parameter inference in these implementation involve numerical integration through MCMC due to the absence of analytical solutions (Tolkoff et al. 2018; Hassler et al. 2022), it can be computationally prohibitive for very large datasets (fitting a model to a dataset with *n* = 1000; *p* = 1000; *q* = 4 takes several hours, see Table S1 in (Hassler et al. 2022)). Furthermore, the current implementation of the PFA is limited to BM and is incompatible with geometric morphometric data processed by Procrustes superimposition techniques, such as the crocodyliform dataset used here, because FA (and consequently PFA) is not rotation-invariant. Although we illustrate the use of the P3CA with geometric morphometric data, it is applicable to other types of datasets, even in fields where the use of ordination approaches is not a regular practice. For instance, gene expression profiles, which typically comprise tens of thousands of traits with missing values, can be analysed with the P3CA to explore the coevolution of genes. The latent variables could reflect functional modules and pathways of co-expressed genes which could be correlated with other traits (e.g., life history traits), shedding light on the biological processes that shape the evolution of gene expression.

### Future directions

Multiple improvements could be envisioned to improve the efficiency of the P3CA and to extend it to more complex models. For example, other evolutionary models based on gaussian processes could be implemented, such as the Ornstein-Uhlenbeck (OU) process with one or multiple optima, or the Early Burst (EB) model. As the P3CA is a probabilistic model with a well-defined likelihood (gaussian distribution), it should be possible to use common model selection criteria, such as the Akaike or Bayesian information criterion, to compare the fit of these models (see model selection approaches proposed in (Caetano and Hearn 2026)). Also, relaxing the assumption that only one evolutionary model underlies all the latent variables would extend the hypotheses that can be evaluated. This would allow to model different set of traits, such as functionally related anatomical structures or groups of interacting genes, with different models of trait evolution (e.g. OU or EB) and/or specific evolutionary parameters. Gu and Shen (2020) recently proposed such a latent variable model that they applied to spatially correlated data. However, this approach is currently limited to complete datasets, and implementing it in the phylogenetic case would be computationally intensive, since it would require the computation of multiple phylogenetic covariance matrices (one for each latent variable) and the optimisation of a larger number of parameters. Further development is required in order to design an efficient EM algorithm to extend the approach from (Gu and Shen 2020) to possibly large and incomplete phylogenetic datasets. Finally, the current implementation of the P3CA could benefit from several improvement to speed up computing time. This is particularly the case when dealing with datasets with missing values, as estimating these values involves many matrix operations that could potentially be replaced by faster (linear time) pruning-like algorithms. Additionally, some methods to accelerate the EM algorithm (Varadhan and Roland 2008; Kuroda and Geng 2023) could be considered in the future.

## Acknowledgments

This work was done thanks to valuable and insightful discussions with Paul Bastide, Philippe Veber, Nils Chabrol, Anaïs Duhamel, Fabien Condamine and Anne-Claire Fabre. The paper also benefited from constructive comments from members of the J.C., H.M. and A.G. teams. We thank the authors of the Crocodyliformes dataset for making their data available, and the computing facilities of the LBBE/PRABI and CC-IN2P3/CNRS that were used to run all our simulations.

## Data Availability Statement

The P3CA is implemented in the function *p3ca()* in the R package ‘mvMORPH’ available on CRAN (https://cran.r-project.org/package=mvMORPH), and GitHub (https://github.com/JClavel/mvMORPH). The geometric morphometric data of the Crocodyliforms skull, the phylogenetic tree and the ecological information used as illustration of the P3CA use can be retrieved from https://github.com/rnfelice/Croc_Skulls.

## Appendix I

### Developing an Expectation – Maximisation (EM) algorithm for the P3CA

We use uppercase letters to refer to matrices and lowercase letter for scalars. *Tr*(***X***), |***X***|, ***X***^−1^, and ***X***^*T*^ indicate the trace, determinant, inverse, transpose of the matrix ***X***, respectively, while ⟨***X***⟩ is its expected value. The matrix dimensions are noted as subscript or specified as (*a* × *b*) for a matrix with *a* rows and *b* columns. ***I***_*a*_ is an identity matrix of dimension *a*, and ⊗ is the Kronecker product. Finally, ***X*** ∼ 𝒩(**Θ, Σ**) means that ***X*** belongs to a multivariate normal distribution with mean **Θ** and variance-covariance **Σ**, while ***X*** ∼ ℳ𝒩_*a,b*_ (**Θ, U, V**) indicates a matrix-variate normal distribution with mean matrix **Θ** (*a* × *b*) and variance-covariance matrices **U** (*a* × *a*) and **V** (*b* × *b*). The matrix variate distribution corresponds to a vectorized matrix with Kronecker structured covariance: *vec*(***X***)∼ 𝒩(*vec*(**Θ**), **V** ⊗ **U**).

In the first part, we describe the mathematical identities used in the next sections. In the second one, we describe the conditional distributions used in the Expectation-Maximisation algorithm, and the formulation for the missing values. Finally, in the third and fourth part, we describe the EM algorithm and the lower-bound of the log-likelihood, respectively.

#### 1. Identities

We use the following identities to develop the EM algorithm:

A. *Expected values* (Gupta and Nagar 2000):
B. *Entropy for a gaussian distribution:* Using the Kronecker identities below, we write the entropy for a matrix variate distribution as:
C. *Other identities for matrices and Kronecker products (Gupta and Nagar 2000):*

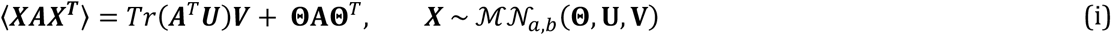

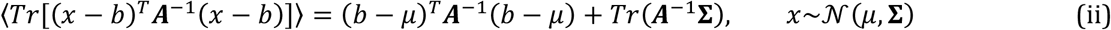

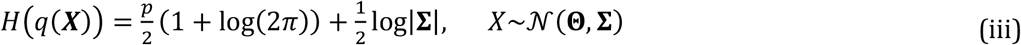

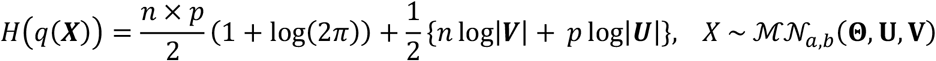

Let A be a *p* × *p* matrix, B a *n* × *n* matrix, and *c* is a scalar.

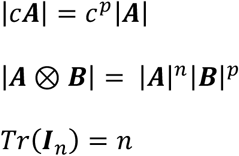

#### 2. Latent variables and missing values

Following the continuous latent variable model in Equation (1) of the main text, ***Y*** is a matrix of traits for *n* species and *p* traits, described by *q* latent variables ***Z*** (*q* × *n*), with parameters ***W*** (*p* × *q*) and *σ*^2^. Hereafter, we use *θ* = {***W***; *σ*^2^} for the last two parameters. The conditional distributions for the observed and latent variables, are defined as:

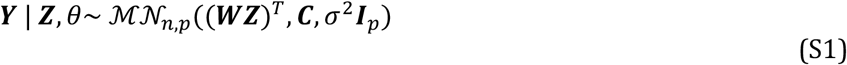

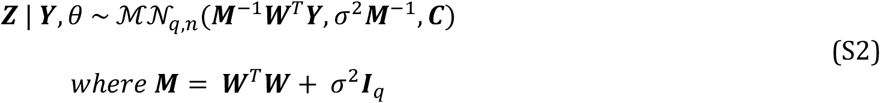

The matrix ***C*** corresponds to the phylogenetic covariance matrix obtained for a given *λ* value, the parameter describing Pagel’s lambda model (***C***^*λ*^).

We note ***Y*** = [*y*(*x*), …, *y*(*x*_*p*_)] with *x* ∈ {1, …, *p*}, and *y*(*x*) = (*y*_1_ (*x*), …, *y*_*n*_ (*x*))^*T*^ the values for the trait *y*(*x*) across *n* species. When there are missing values in ***Y***, the data can be partitioned into observed (***Y***_[*o*]_) and missing values (***Y***_[*m*]_):

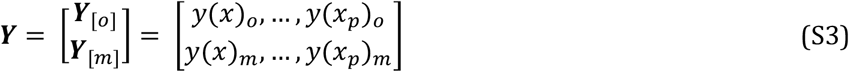

Conditional on the observed values (noted *y*(*x*)_*o*_), the latent variables and the parameters *θ*, the missing values in column *x* of ***Y***, noted *y*(*x*)_*m*_, are assumed to be normally distributed random variables. The conditional distribution for the missing values is given by:

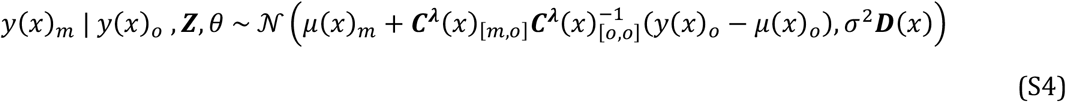

Where,

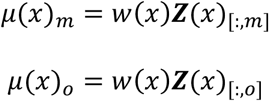

And

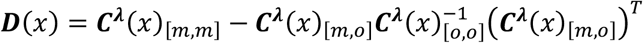

The vector *w*(*x*) contains the *q* latent variables for the trait *x* (i.e., the row of ***W*** corresponding to trait x), and ***C***^*λ*^(*x*) is the matrix with the phylogenetic covariances between missing values (***C***^*λ*^(*x*)_[*m,m*]_), between missing and observed values (***C***^*λ*^(*x*)_[*m,o*]_), or between observed values (***C***^*λ*^(*x*)_[*o,o*]_) for the trait *x*. Thus, the *q* latent variables ***Z*** are partitioned according to the observed (***Z***(*x*)_[:,*o*]_) and missing (***Z***(*x*)_[:,*m*]_) values for the trait *y*(*x*). For both ***Z***(*x*) and ***C***^*λ*^(*x*), the subscripts indicate rows and columns positions for which species are missing (m) or observed (o) for the trait *y*(*x*) in the complete matrices.

#### 3. An Expectation-maximisation (EM) algorithm for the P3CA

To develop an EM algorithm to infer the parameters describing the continuous latent variable model of the P3CA, conditional of a given *λ* value (i.e., the phylogenetic covariance matrix under the Pagel’s lambda model), we should first define the complete-data log-likelihood, or joint log-likelihood of both ***Y*** and ***Z***. Using the conditional distributions (equations 2 and 3 in the main text), the joint log-likelihood is:

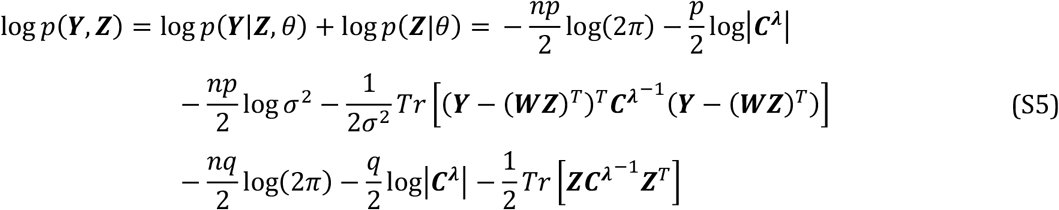

In the E-step of the EM, the expectation of the joint log-likelihood is taken with respect to the conditional distributions of both the latent variable and the missing values for trait *y*(*x*), *q*(***Z***) and *q*(*y*(*x*)_*m*_), respectively. These distributions are:

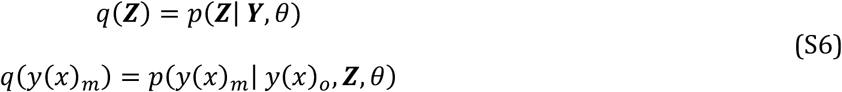

Applying the identity (i), the expectation for log *p*(***Y, Z***) with respect to *q*(***Z***) is – for the terms involving ***Z*** (see also(Li et al. 2009)):

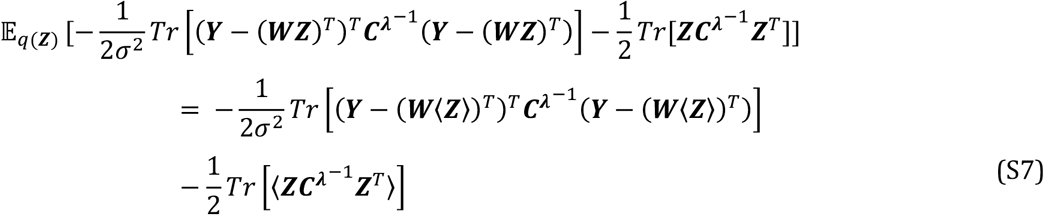

Where,

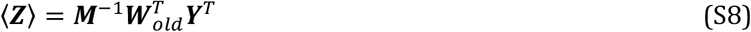

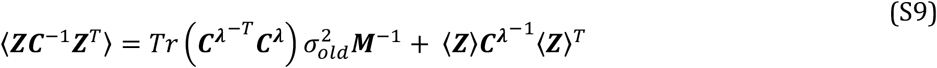

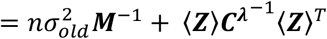

and,

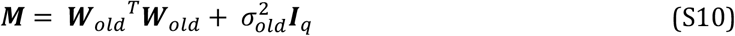

Similarly, applying the identity (ii), the expectation for log *p*(***Y, Z***) with respect to *q*(***Y***_[*m*]_), and only considering the terms involved for the missing values in ***Y***, is given by:

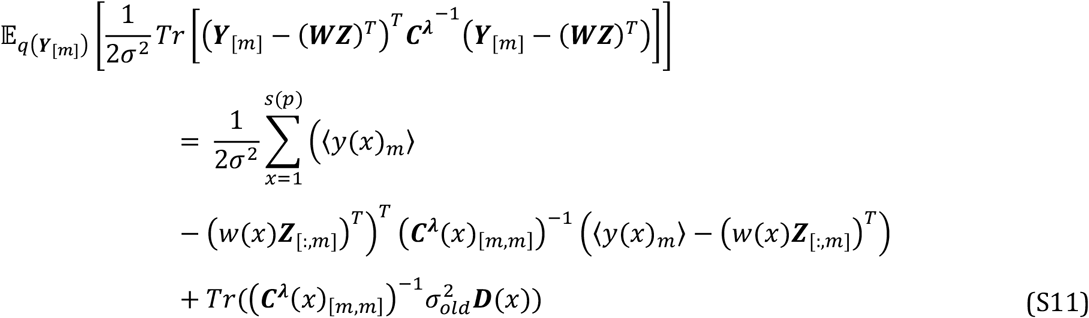

Where,

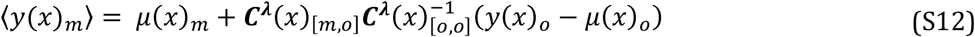

Where *s*(*p*) is the set across the *p* traits containing missing values. The integration with respect to *q*(***Y***_[*m*]_) of the quadratic form in log *p*(***Y, Z***), show that the expectation (eq. S12) is obtained by replacing the missing values in ***Y*** by the conditional mean ⟨*y*(*x*)_*m*_⟩ for each trait and by adding a term related to the phylogenetic covariances. After some manipulations, the equation S11 can be expressed as:

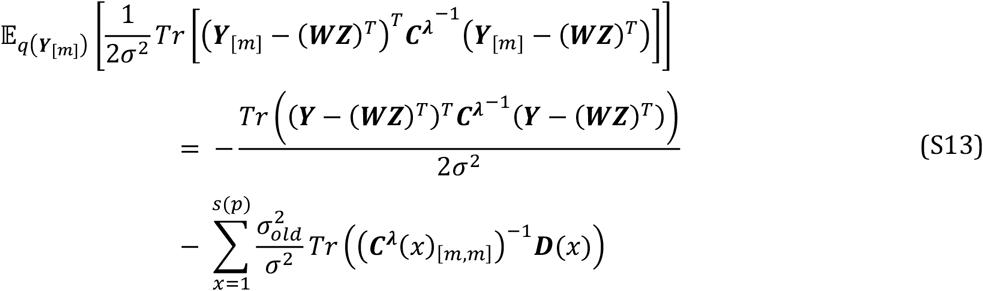

Where the missing values in ***Y*** were filled in by the conditional expectations for each trait (i.e., ⟨*y*(*x*)_*m*_⟩). Combining the expectations for the terms in the joint log-likelihood with respect to the distribution of the missing values and the latent variables, we obtain the expectation of the complete-data (joint) log-likelihood as:

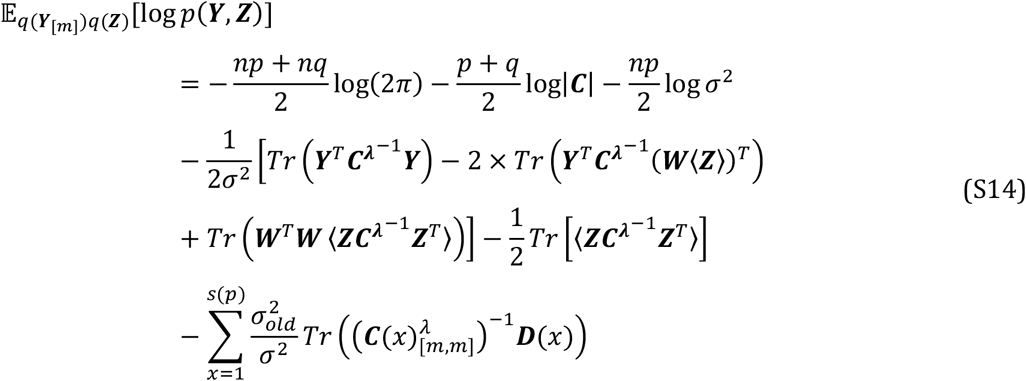

Using equation S14, we derive the solutions for the parameters ***W*** and *σ*^2^ as follow:

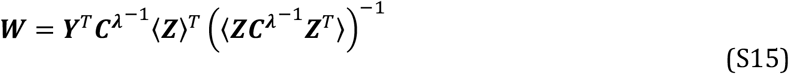

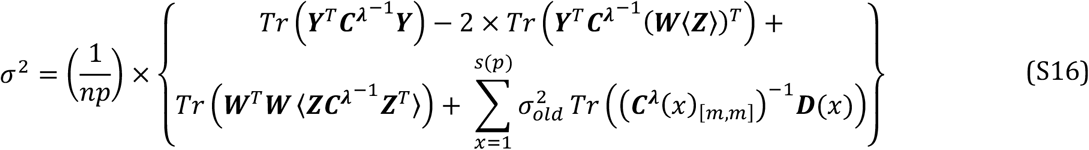

The EM algorithm iterates until convergence between the *E-step*: estimating the expectation of the joint log-likelihood with respect to *q*(*Z*) (equations S8 and S9) and *q*(***Y***_[*m*]_) (equation S11) based on current parameters estimate; and the *M-step*: estimating new ***W*** and *σ*^2^ parameters that maximise the expected joint likelihood (equations S15 and S16).

#### 4. Lower-bound of the log-likelihood

Finally, we obtain the lower-bound for the log-likelihood of ***Y*** (*p*(***Y***)) by adding the entropy for *q*(***Z***) and *q*(***Y***_[*m*]_) to the expected log-likelihood (Roweis and Ghahramani 1999).

Using the identity (iii), we can define the differential entropy for *q*(***Z***) and *q*(*y*(*x*)_*m*_) as:

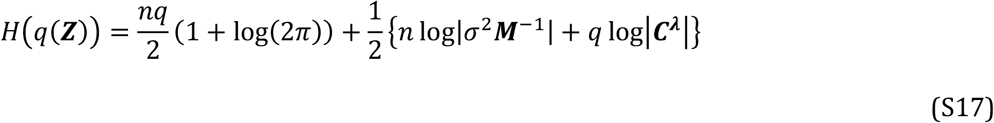

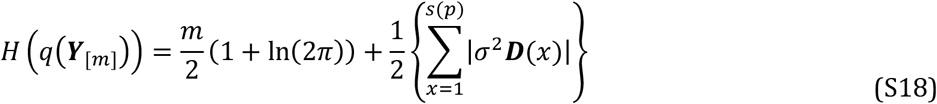

Where *m* is the total number of missing values in the dataset.

Adding the entropy terms in S17 and S18 to the expectation of the complete-data log-likelihood (S14), we obtain the lower-bound for the log-likelihood of *p*(***Y***) as:

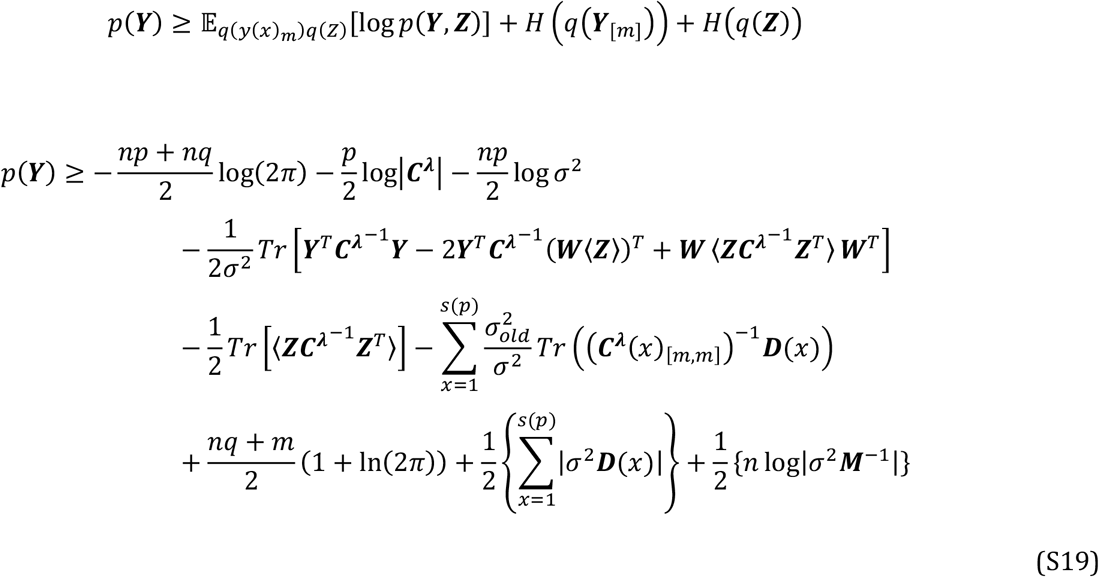

We maximise the lower-bound of *p*(*Y*) (S19) to infer both, the parameters of the continuous latent variable model, and the parameter *λ* for the Pagel’s lambda model.

## Appendix II

### Implementation of the EM algorithm for the P3CA and optimisation of Pagel’s lambda parameter

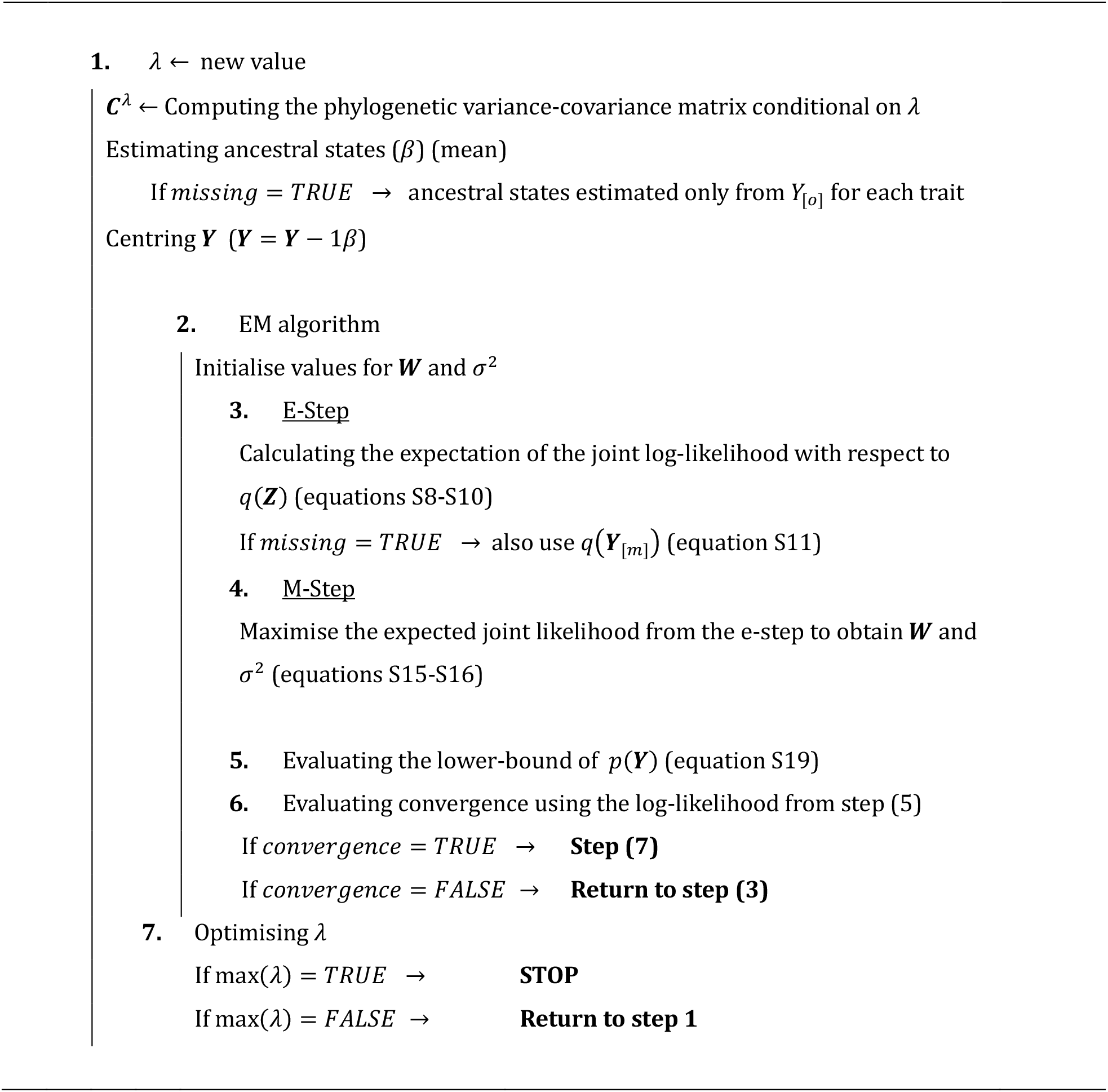

## Appendix III

### Efficient estimation of the marginal log-likelihood for the P3CA

Although the marginal log-likelihood (equation 5 in the main text) includes some potentially large matrices (*p* × *p*), we take advantage of the low-rank structure of the latent variable model for estimating it more efficiently. The main bottleneck in equation 5 is the trace term *Tr*((***WW***^*T*^ + *σ*^2^***I***_*p*_)^−1^ (***Y*** − **Θ**)^*T*^***C***^−1^(***Y*** − **Θ**)) and the log-determinant log|(***WW***^*T*^ + *σ*^2^***I***_*p*_)|, which both involve *p* × *p* matrices.

Estimating 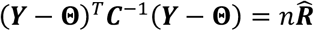 efficiently is possible by applying a pruning algorithm on the phylogenetic tree for obtaining the inverse of the square root of the matrix ***C*** 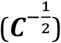, and multiplying it to the centred trait matrix 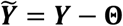:

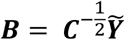

This operation is computationally cheaper than explicitly inverting the covariance matrix ***C***, and is equivalent to computing the phylogenetic independent contrasts (see (Stone 2011)). Using the above transformation, the trace term can be expressed as *Tr*((***WW***^*T*^ + *σ*^2^***I***_*p*_)^−1^***B***^*T*^***B***). We can efficiently estimate this term using the singular value decomposition (SVD) 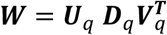 rather than by estimating the term (***WW***^*T*^ + *σ*^2^***I***_*p*_)^−1^ explicitly. To exploit this, we note that the trace term in the log-likelihood can be decomposed as:

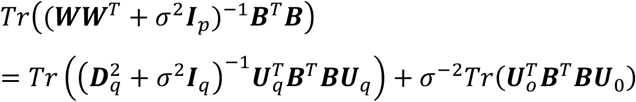

Where ***U***_*q*_ correspond to the left singular vectors corresponding to the *q* non-null singular values 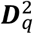 of ***WW***^*T*^, while ***U***_0_ are the *p-q* singular vectors of the null complement (corresponding to the *p-q* null singular values). Taking the projection of the centred data ***B*** onto the subspace spanned by ***W*** using the singular vectors matrix ***U***_*q*_ we can obtain the first term in the right-hand side of the equation above as:

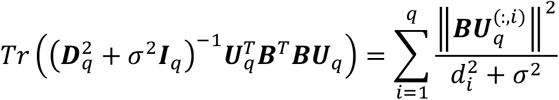

The operator ‖·‖^2^ = ∑|*i*|^2^ is the squared norm of a matrix, and ***U***^(:,1)^ corresponds to the *i* column of the matrix ***U***. To obtain the null complement (i.e., the space corresponding to the *p-q* singular values and singular vectors of the matrix ***WW***^*T*^ + *σ*^2^***I***_*p*_), we can take the orthogonal projection of ***B*** onto the complement of the column space spanned by ***W*** (i.e. 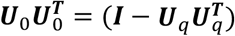):

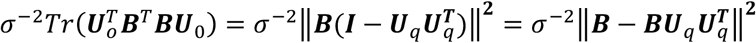

Where ***BU***_*q*_ was already computed for the first term. The SVD of a *p* × *q* matrix can be estimated more efficiently than the inverse (or SVD) of a *p* × *p* matrix (Ilin et al. 2010; Golub and Van Loan 2013). Decomposing the trace term and using the SVD of ***W*** enables us to employ sums of squared terms in smaller matrices, a process that can be carried out very efficiently. Substantial computational savings can be achieved when *q* ≪ *p*.

Likewise, taking advantage of the singular value decomposition of ***W***, we can compute efficiently the log-determinant term log| (***WW***^*T*^ + *σ*^2^***I***_*p*_)| as:

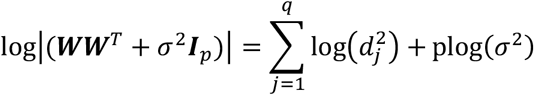

This formulation prevents the computation of large matrices, focusing on simpler and less computationally demanding operations, making the approach scalable to very high-dimensional problems when there are no missing values in the dataset.

