## Supplementary material for "A probabilistic and phylogenetic principal component analysis for modelling high-dimensional trait evolution"

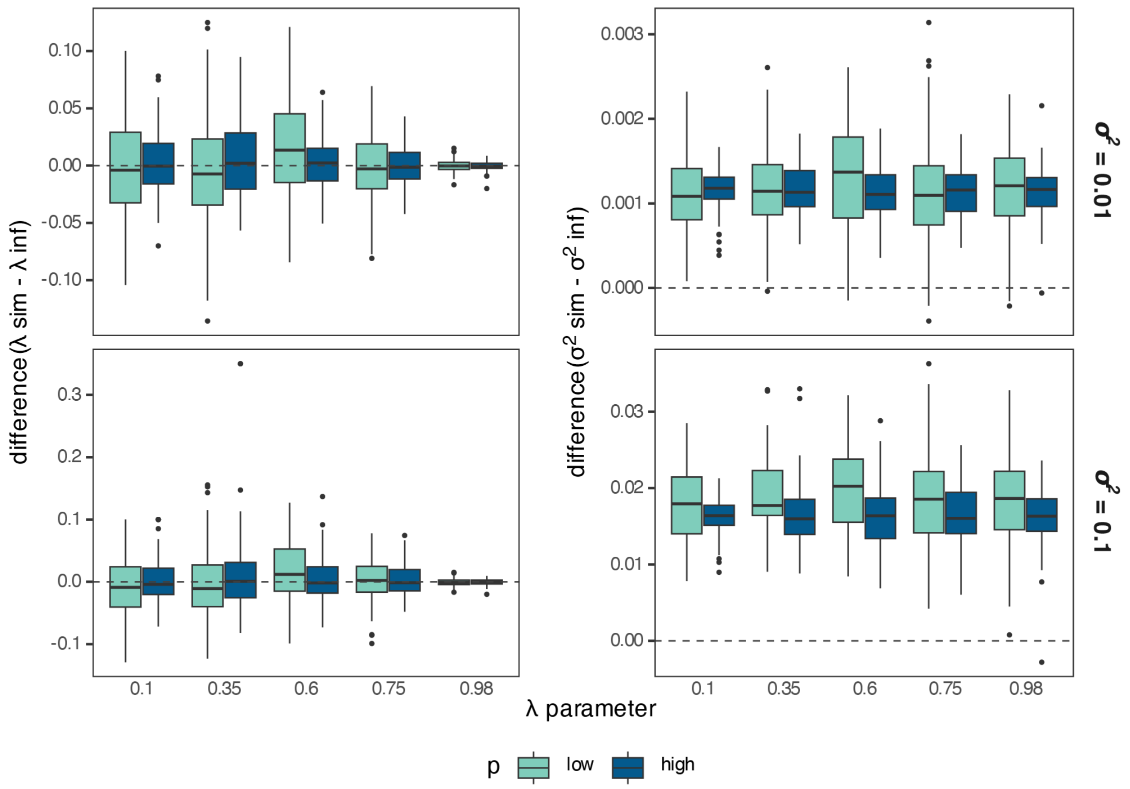


Figure S1. Parameter inference for $\lambda$ and $\sigma^2$ across different $\lambda$ values, using the analytical solutions for $\sigma^2$and $\boldsymbol{W}$ (equation 6 in main text). The plots show the difference between the simulated and the inferred parameter. The parameter $\lambda$ is well estimated across different values with higher precision at larger values (stronger signal). The parameter $\sigma^2$ is slightly underestimated although this bias does not increase or decrease with the $\lambda$ value. Results are shown for 100 simulations (from 1st to 3rd quantile in boxes, while median in solid line) with $n = 50, q = 5, \sigma^2 = 0.1$, and $p = 25$ (low-dimension) or $p = 100$ (high-dimension).

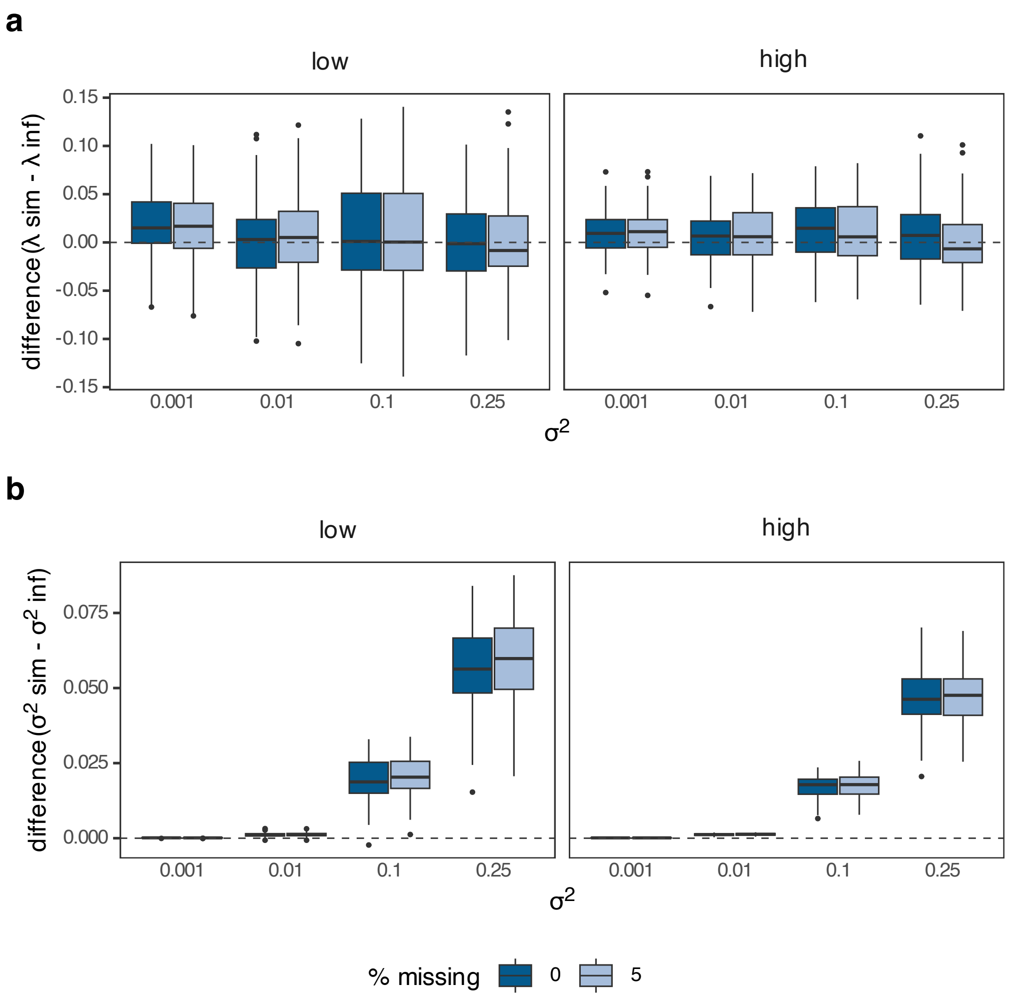


Figure S2. Parameter inference for $\lambda$ and $\sigma^2$ across different values for $\sigma^2$, for complete and incomplete (5% of missing values) datasets. The inference is made with the EM algorithm. While the inference for $\lambda$ is almost unbiased, $\sigma^2$ is underestimated when the simulated values of $\sigma^2$ are large. No differences are detected between low and high-dimensional conditions. The plots show the results (from 1st to 3rd quantile in boxes, while median in solid line) across 100 simulations with $n = 50, q = 5, \sigma^2 = 0.1$, and $p = 25$ (low-dimension) or $p = 100$ (high-dimension).

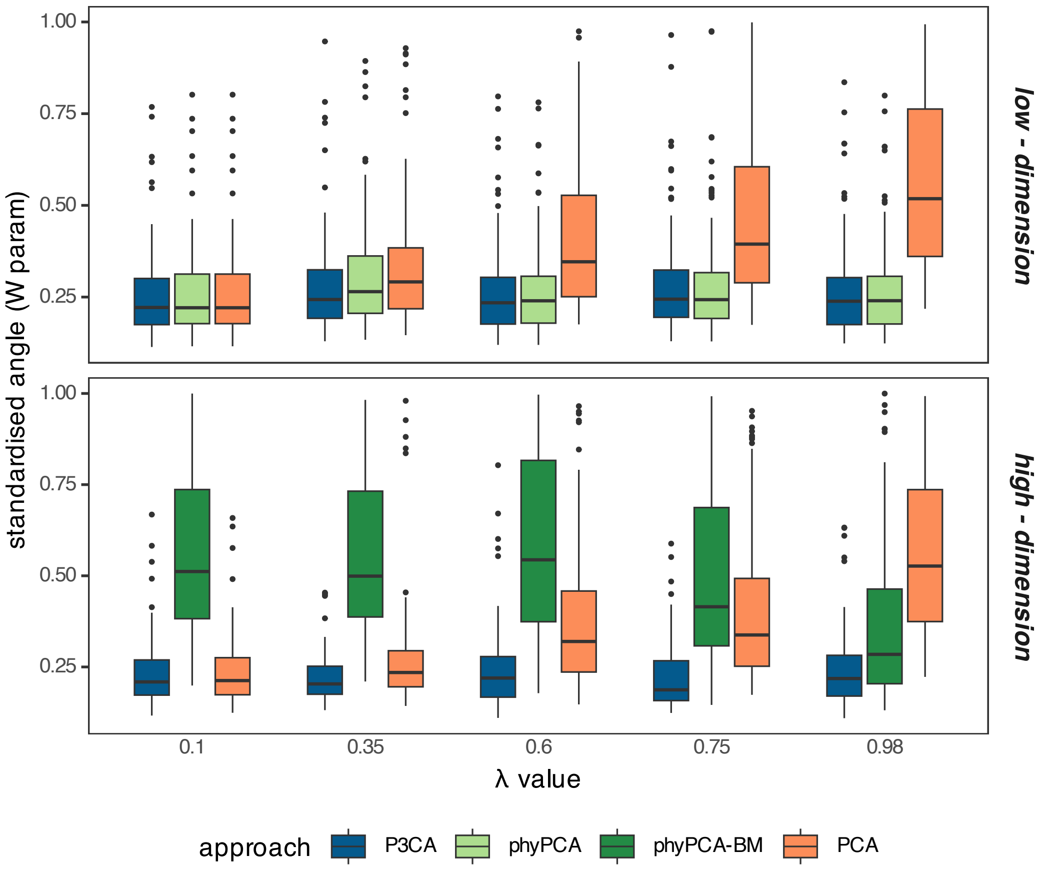


Figure S3. Parameter inference for $\boldsymbol{W}$ using analytical solution (equation 6 in main text) across different $\lambda$ values. The $\boldsymbol{W}$ matrix is compared to the simulated matrix using the last standardised principal angle between their subspaces. Comparisons to the eigenvectors obtained from the phylogenetic PCA under Pagel's lambda model (phyPCA) and Brownian-motion (phyPCA-BM), and the conventional PCA are also shown. The smaller the standardised angle between the inferred and the simulated $\boldsymbol{W}$matrices, the closer are the matrices. Unlike other approaches, inference of $\boldsymbol{W}$ from the P3CA is unaffected by the value of $\lambda$ and performs best across most conditions. In high-dimensions, Brownian-motion is the only available model for phyPCA. Resutls are shown across 100 simulations (from 1$^{st}$ to 3$^{rd}$ quantile in boxes, while median in solid line) with $n = 50, q = 5, \sigma^2 = 0.1$, and $p = 25$ (low-dimension) or $p = 100$ (high-dimension).

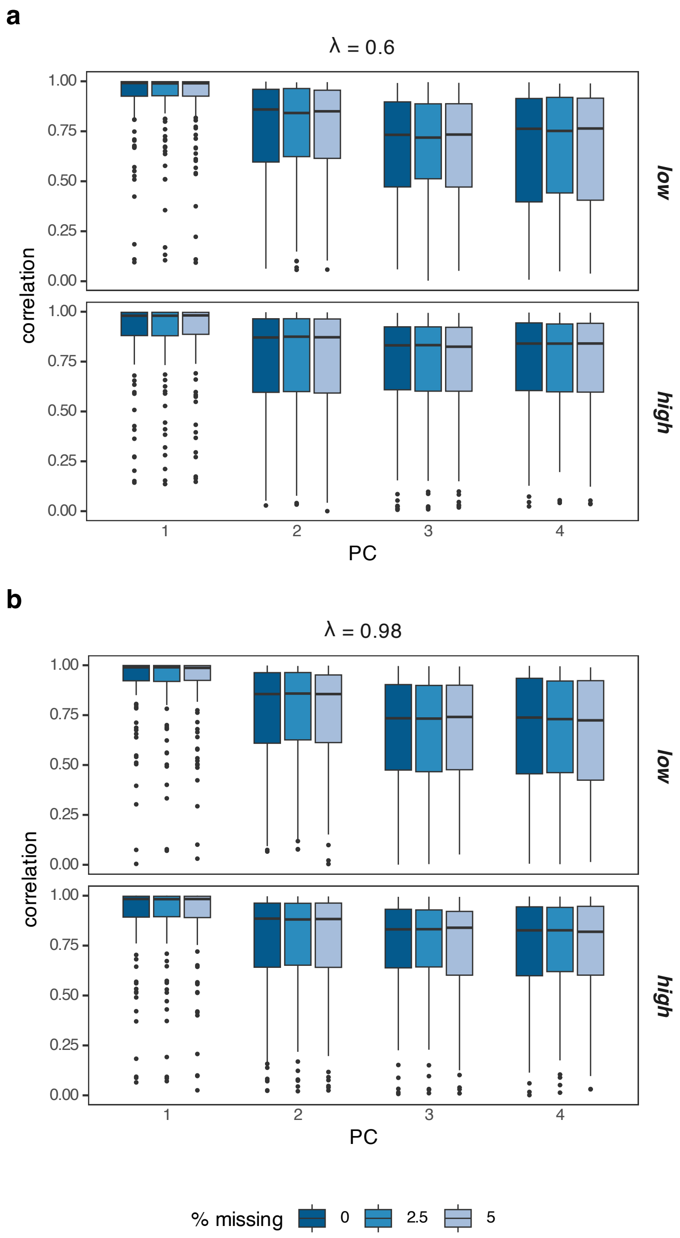


Figure S4. Correlation between the principal components (PC) obtained using the simulated parameters and those inferred from the P3CA with the EM algorithm. The correlation does not change with either the proportion of missing values, the dimensionality (low or high-dimension) or the value for $\lambda$ (Brownian motion model when $\lambda = 1$). The correlation decreases as the variance explained by the component decreases (lower variance in the four component). The plot shows the correlations (from 1st to 3rd quantile in boxes, while median in solid line) across 100 simulations with $n = 50, q = 5, \sigma^2 = 0.1$, and $p = 25$ (low-dimension) or $p = 100$ (high-dimension).

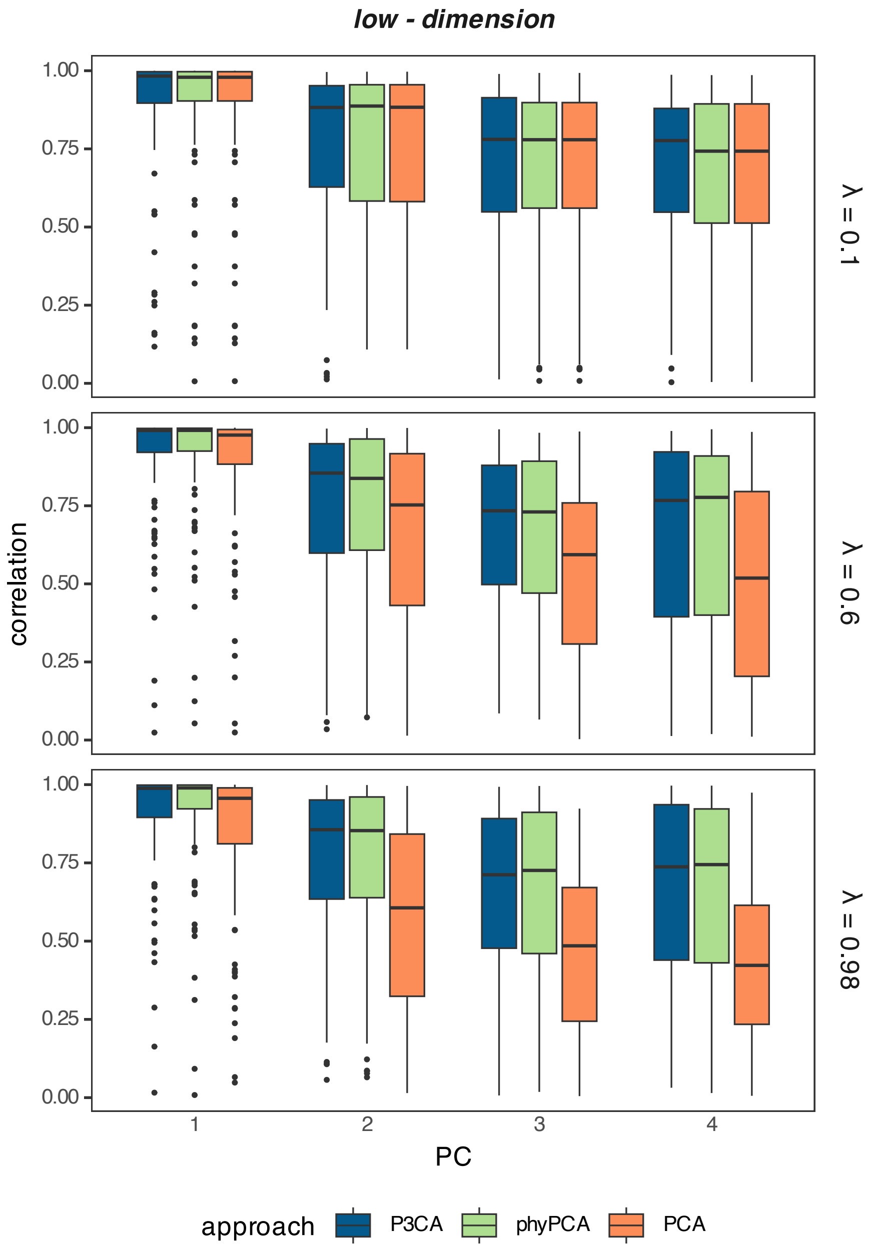


Figure S5. Correlation between the principal components (PC) obtained using the simulated parameters and those inferred by the P3CA, the conventional PCA and the phylogenetic PCA (phyPCA) with the EM algorithm, in low-dimensional settings. The phylogenetic PCA (phyPCA) is fitted under Pagel's lambda model. The P3CA and phyPCA perform the best (higher correlation) while, as expected, the performance of the conventional PCA decreases with the $\lambda$ value. The correlations are shown across 100 simulations with $n = 50, q = 5, \sigma^2 = 0.1$, and $p = 25$ (low-dimension).

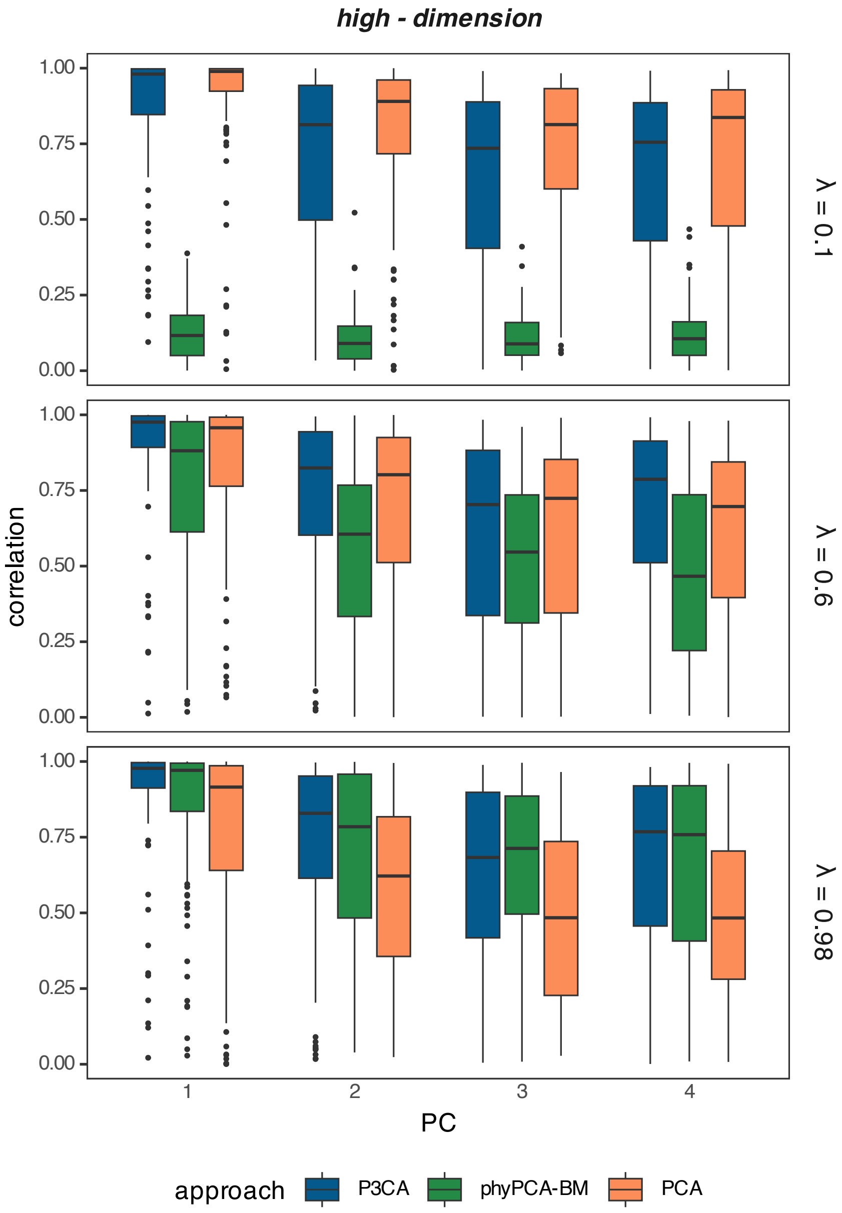


Figure S6. Correlation between the principal components (PC) obtained using the simulated parameters and those inferred by the P3CA, the conventional PCA and the phylogenetic PCA with the EM algorithm, in high-dimensional settings. The phylogenetic PCA (phyPCA) is fitted under the Brownian-motion model (phyPCA-BM), as this is the only model applicable in high-dimensional data. The P3CA consistently performs as the best or one of the best across most conditions. In contrast, the performance of the PCA and phyPCA-BM depends on the value of $\lambda$. The plots show the correlations (from 1st to 3rd quantile in boxes, while median in solid line) across 100 simulations with $n = 50, q = 5, \sigma^2 = 0.1$, and $p = 100$ (high-dimension).

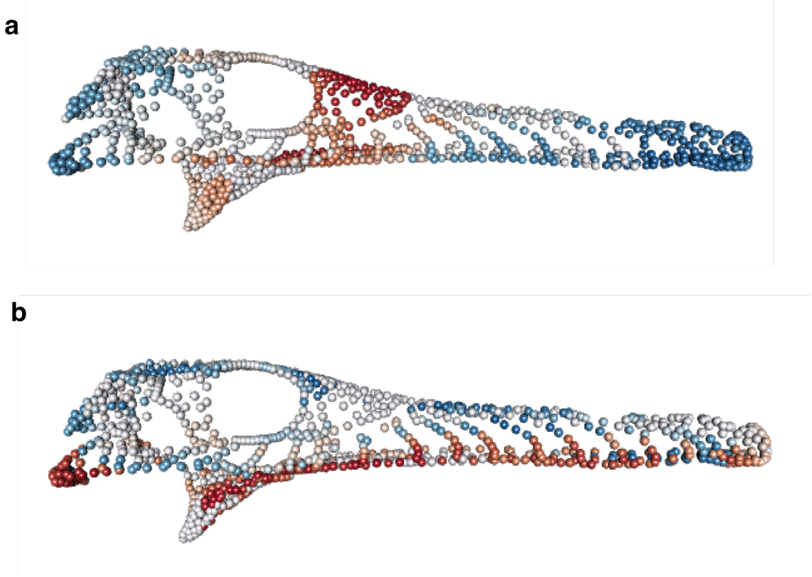


Figure S7. Localisation of the landmarks and semi-landmarks of crododyliformes skulls with the highest variation along the first component obtained by the P3CA, under the **a**) Pagel's lambda model, and the **b**) Brownian motion model. The colours represent the landmark or semi-landmark with coordinates having the highest PC loading on PC1 (while blue colours indicate a negative loading value, red colours indicate a positive one). The loadings are the correlations between the principal component and each coordinate (from either the landmarks or semi-landmarks). The stronger the loading value, the greater is its contribution to the component. To make it easier to visualise, only the landmarks and semi-landmarks with loadings greater than 0.2 or less than -0.2 in at least one of their axes (x, y or z), are coloured. The first component mostly explains changes in similar regions for both models except for the frontal and prefrontal regions. For these two anatomical regions, changes on PC1 are only observed when the P3CA was fitted with the Pagel's lambda model. The figure shows the projection of the landmarks and semi-landmarks onto the same mean shape.

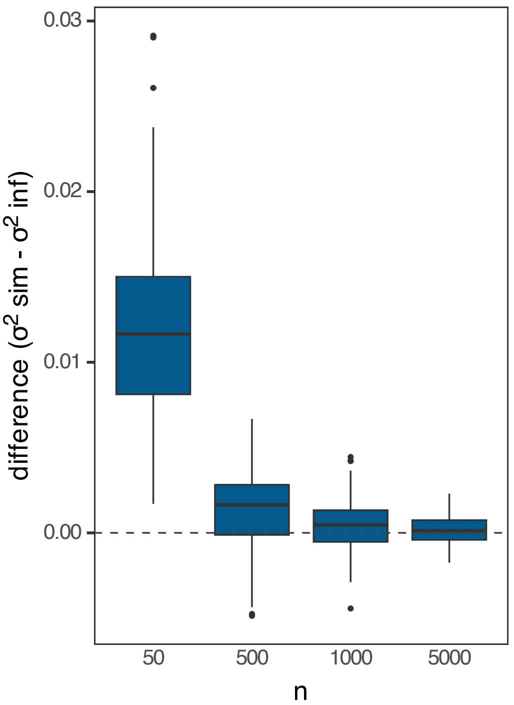


Figure S8. Difference between the simulated $\sigma^2$ parameter and the inferred one for different $n$ values, when $p = 25$. The underestimation for $\sigma^2$ decreases as $n$ increases. The plot shows the results for 100 simulations with $q = 5$ and $\sigma^2 = 0.1$. In the boxplots, boxes include from 1st to 3rd quantile, while median is shown in solid line.

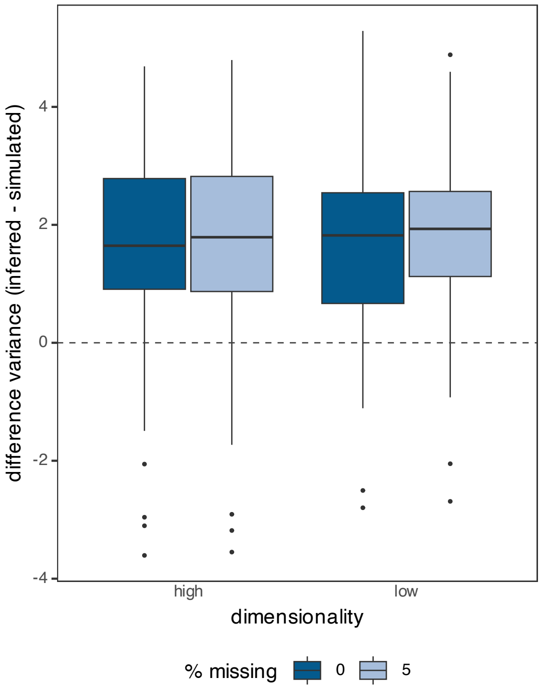


Figure S9. Difference variance explained between the inferred and simulated lower-dimensional space. The difference is shown for low and high-dimensional conditions with complete and incomplete (with 5% of missing values) datasets. The explained variance is overestimated, which is consistent with the underestimation of $\sigma^2$ (see *Parameter Inference* in the *Discussion* section of the main text). The plot shows the difference across 100 simulations with $n = 50, q = 5, \sigma^2 = 0.1,$ and $p = 25$ (low-dimension) or $p = 100$ (high-dimension). In the boxplots, boxes include from 1st to 3rd quantile, while median is shown in solid line.

*Exploring the morphospace for the skull shape in Crocodyliformes*

The differences in morphospaces obtained with the P3CA under Pagel's lambda and Brownian motion persist in the third and fourth components.

With Pagel's lambda model, the third component is principally explaining variation around the frontal, nasal and premaxilla bones, describing skulls with a concave snout (for example, the extreme *Simosuchus clarki*) to slightly convex skulls (see for example *Crocodylus palustris*). The pterygoid position is also changing, moving from a vertical to a more horizontally and backward-facing position. The fourth component reinforces changes observed in the pterygoid, but also show some changes around the frontal and postorbital regions that represent a shift from a lateral orbit to a more dorsal one. PC3 and PC4 cluster the fresh-water lineages while the terrestrial omnivorous and marine lineages are dispersed in their periphery. Two morphologies stand out as outliers: the presumably terrestrial *Simosuchus clarki* and the extinct *Stangerochampsa mccabei* (Figure S10a-b). This pattern reflects the distinct morphology of *Simosuchus clarki* in our sample, with a short and dorsoventral enlarged skull. The case of *Stangerochampsa mccabei* is less clear, as its morphology is more similar to that of extant Crocodyliformes. However, the distance to the main group may suggest specific features around the pterygoid and the postorbital regions. The distance between these two lineages and the rest of the group may be due to the small sample size (then, increasing the sample size across all known morphologies is needed). Alternatively, it may indicate different evolutionary processes underlying the evolution of the anatomical parts involved in the axes. In contrast, when the P3CA is performed with a Brownian-motion model, the third and fourth components exhibit no ecological patterns. The changes in the skull are less perceptible with the main variation in the third component, concentrated around the maxilla, quadrate and quadratojugal regions. Along the fourth, the regions with the higher variation are sparse around the skull with some around the orbits, the nasal region, the quadrate, the condyle and slightly in the pterygoid (Figure S10c-d). All the inferences are based in the phylogenetic tree shown in figure S11.

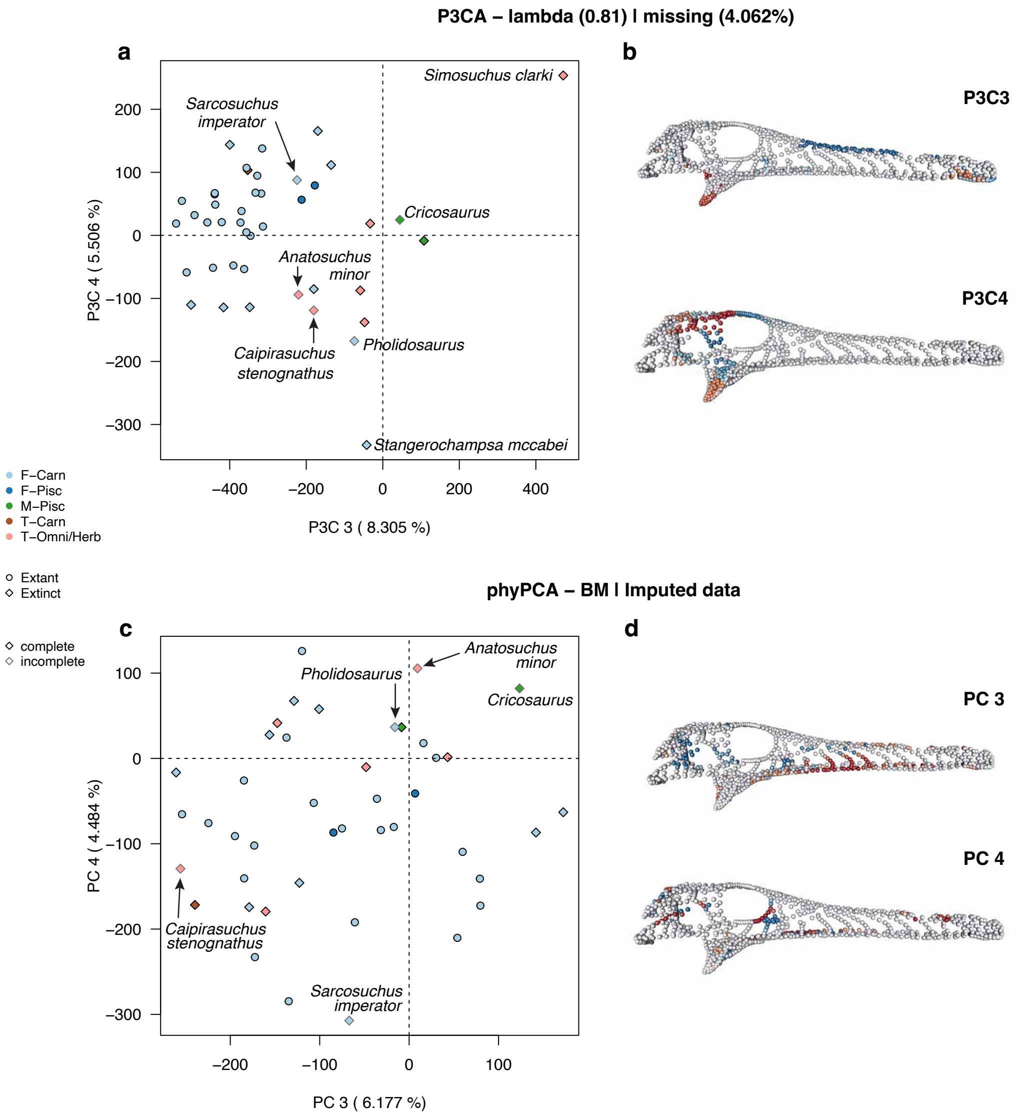


Figure S10. Morphological space of skull shape in Crocodyliformes is shown for the third and fourth principal components of the **a**) P3CA (top panel) and **b**) phylogenetic PCA under Brownian-motion model (phyPCA-BM, bottom panel). Only species with missing values are named in the morphospace. The figure also shows the anatomical changes (landmarks with the greatest variation) along the third and fourth component under the **b**) Pagel's lambda model, and the **d**) Brownian motion model. The colours represent the highest or lowest loading value across the three coordinates describing each landmark or semi-landmark. As for previous figures, only landmarks and semi-landmarks with loadings greater than 0.2 (cold colours) or lesser than -0.2 (warm colours), in at least one of their coordinates (x, y or z), are coloured to make visualisation easier. The stronger the loading value, the greater the contribution of the corresponding coordinate to the component. The figures show the projection of the landmarks and semi-landmarks onto a mean shape.

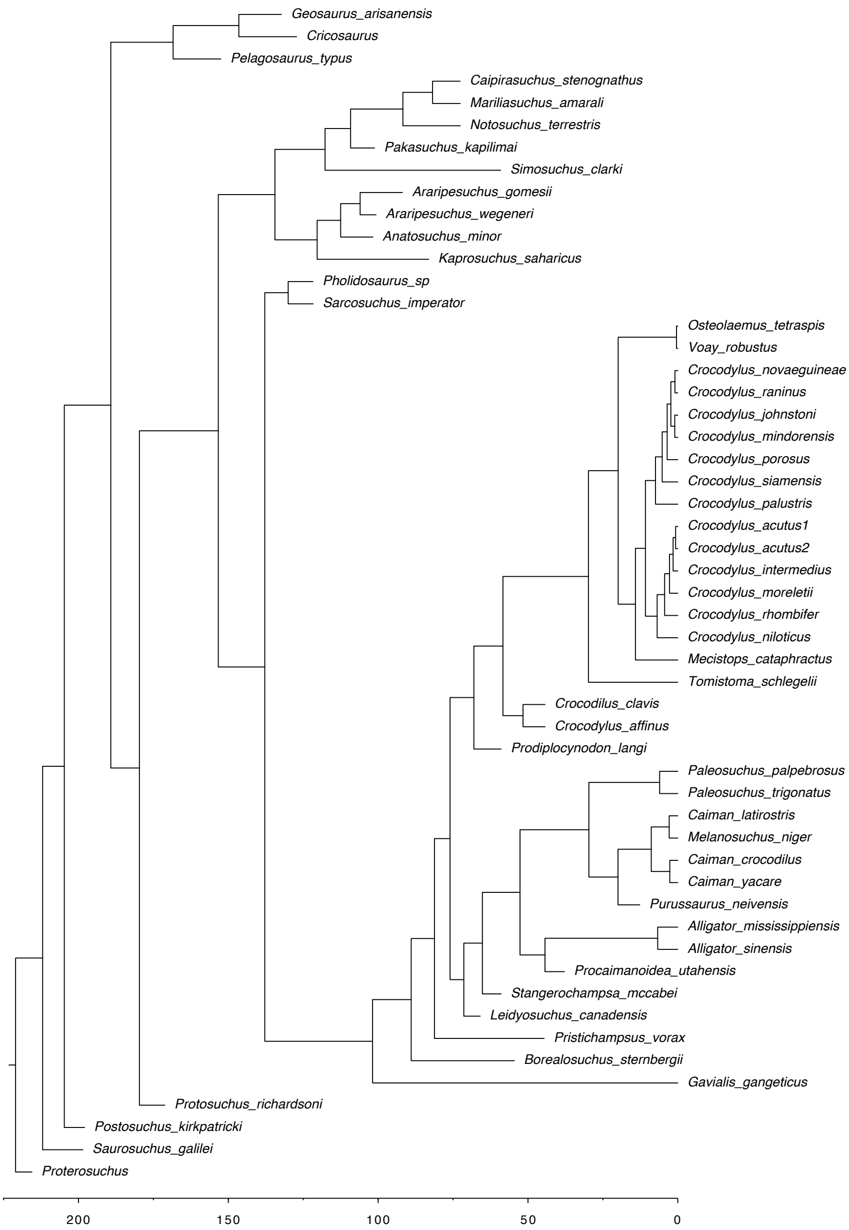


Figure S11. Phylogenetic tree of Crocodyliformes used for fitting the models with the P3CA and the phylogenetic PCA. The time scale is shown on the X-axis. The phylogenetic tree was obtained from (Felice et al. 2021).
